# Optimized large-scale longitudinal biorepository of gastroesophageal adenocarcinoma patient-derived organoids: High-fidelity models for personalized treatment to overcome resistance

**DOI:** 10.1101/2025.07.09.663874

**Authors:** Mingyang Kong, Sanjima Pal, Shuyuan Wang, Julie Bérubé, Ruoyu Ma, Yifei Yan, Wotan Zeng, France Bourdeau, Betty Giannias, Hong Zhao, Nathan Osman, Kulsum Tai, Hellen Kuasne, James Tankel, Greta Everisto, Pierre Fiset, Xin Su, Swneke Bailey, Morag Park, Nicholas Bertos, Veena Sangwan, Lorenzo Ferri

**Affiliations:** Department of Surgical and Interventional Sciences, McGill University, Montreal, Canada; Cancer Research Program, Research Institute of the McGill University Health Centre, Montreal, Canada; Department of Surgery, McGill University, Montreal, Canada; Goodman Cancer Institute, McGill University, Montreal, Canada; Division of Thoracic and Upper Gastrointestinal Surgery, McGill University Health Centre, Montreal, Canada; Department of Pathology, McGill University, Montreal, Canada; Department of Biochemistry, McGill University, Montreal, Canada; Department of Oncology, McGill University, Montreal, Canada

**Keywords:** Biobanking, Patient-derived Organoids, Cancer Model, Precision Oncology, Highthroughput drug Screening, Esophageal Cancer, Gastric Cancer

## Abstract

A major limitation in studying gastroesophageal adenocarcinoma (GEA) has been the lack of reliable models that represent the disease’s complexity. We present lessons learned from a comprehensive large-scale biobanking effort combining traditional sample collection with several *in vitro* models including 3-dimensional patient-derived organoids (PDOs), 2-dimensional cancer-associated fibroblasts (CAFs), tumor-infiltrating lymphocytes (TILs) and/or *in vivo* xenografts. This initiative started in 2018, integrating multiple advanced *ex-vivo* models such as PDOs, patient-derived xenografts (PDXs) and organoids (PDXOs). This unique resource now includes tumor avatars from over 380 consented patients, making it the largest living GEA biobank in the world. We achieved > 90% success rate in creating per-patient models, including 227 tumor-derived and 203 neighboring normal PDOs. These organoids accurately mirror key features of the original tumors, such as their histology (e.g. microsatellite instability), mutations, and drug response, across treatment points. Notably, PDOs can predict individual patient responses to chemotherapy within five weeks, underscoring their clinical relevance. Furthermore, high-throughput drug screening on PDO subsets generates personalized chemosensitivity profiles for 22 drugs. Through a process of continued refinement of culture techniques and tumor sampling approach, our large-scale comprehensive collection of GEA avatars represents a unique and valuable preclinical experimental resource for precision oncology.

**Graphical abstract:** Schematic depiction of GEA live-banking workflow

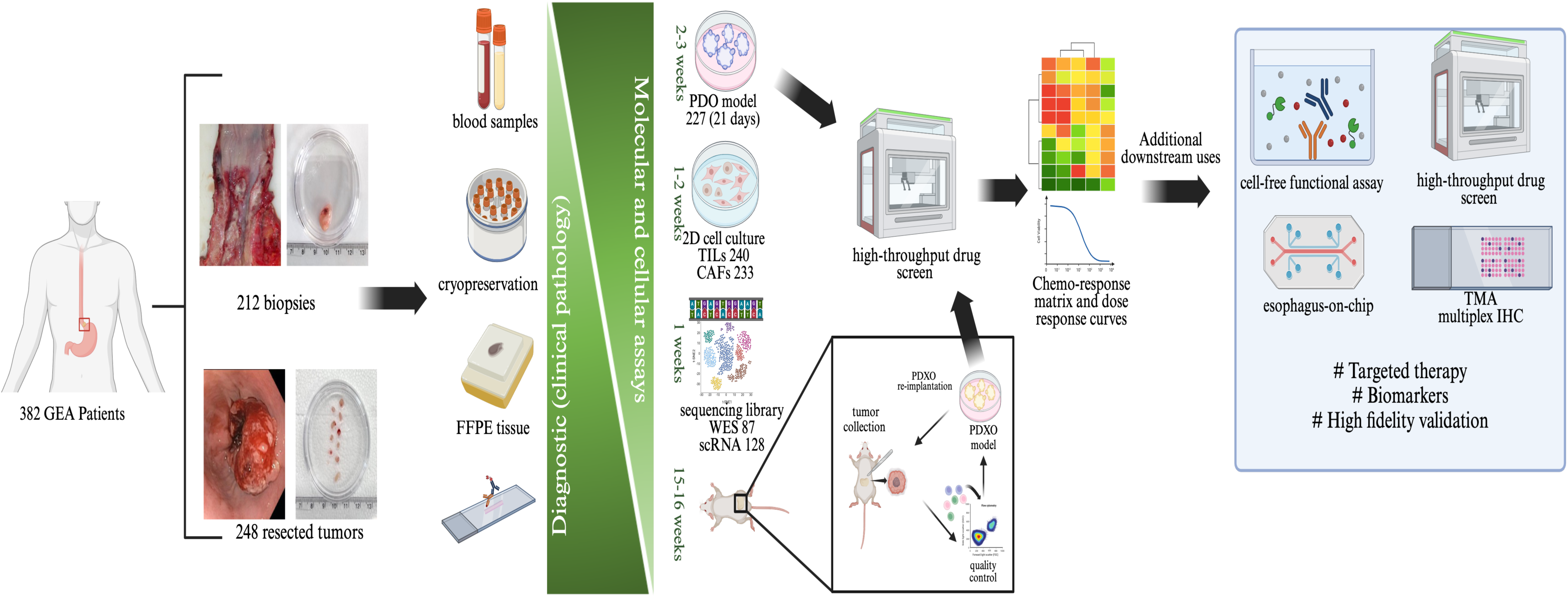

## 1. Introduction

Stomach and esophageal cancers rank as the fifth and seventh causes of cancer-related deaths globally, with over 1.4 million incidences and 1.1 million deaths ^[1–2]^. In the absence of early symptoms and widespread screening, most GEA patients are only diagnosed when symptomatic, generally corresponding to at least locally advanced disease ^[3]^. This, combined with a paucity of biomarker-based or personalized systemic therapies, drives an overall poor prognosis with a 5-year survival of ca. 20% ^[4]^. Although GEA is regarded as a single disease entity based on genomic analysis, these tumors display exceptionally high intra- and inter-tumoral heterogeneity, which likely underlies resistance to standardized systemic therapies ^[5–6]^. Cytotoxic taxane-based perioperative chemotherapy (e.g., FLOT: 5-Fluorouracil, Leucovorin, Oxaliplatin, Docetaxel/Taxotere or DCF: Docetaxel/Taxotere, Cisplatin, and 5-Fluorouracil) before (neoadjuvant) and after (adjuvant) surgical resection with or without concomitant radiotherapy represents the only current standard-of-care (SOC) treatment for locally advanced GEA ^[7]^. However, around 40% of patients display innate resistance to chemotherapy in the neoadjuvant setting, and half of initial responders, as well as many non- or poor responders, later go on to metastatic disease ^[8]^.

Targeted therapies (e.g., anti-HER2 approaches) have shown only limited success ^[9]^, and there are few potential targets with widespread incidence. While clinical trials investigating immunotherapy (primarily targeting PD-1/PD-L1) show some promise, the benefits appear to be either independent of PD-L1 expression in tumor cells or occur regardless of PD-L1 combined positive score (CPS) ^[10–13]^. Thus, there are unmet needs both for the identification of patients unlikely to respond to current SOC therapy in the neoadjuvant setting, and for discovery, validation and establishment of biomarkers and associated novel therapeutic options to address non-responders.

Faithful preclinical models to predict response to SOC therapy or test potential alternative approaches in a clinically useful timeframe are limited. Traditional reductive 2D cell models inadequately recapitulate cancer features such as therapy resistance and hypoxia due to the altered growth environment ^[14–16]^. Rodent-based xenografts require long lead times for generation and cannot easily be scaled up for high-throughput assays; thus, they are not suitable for functional drug screening or real-time clinical decision-making ^[17]^. For example, a large-scale gastric cancer project using solely PDX models has been previously reported ^[18]^. Although the success rate was noteworthy (100 models generated from 349 patients; 29%), the mean initial latency was 74 days, combined with the one-mouse-one-treatment capacity, these factors render it impossible to use this approach for drug screening to direct clinical decision-making in real-time. Patient-derived organoid (PDO)-based approaches promise to act as rapidly accessible higher-fidelity models for drug testing ^[19–22]^. With the amendment of the FDA Modernization Act 2.0 in December 2022, organoids are now permitted as models for drug screening and sensitivity evaluation, offering a faster, high-fidelity and high-throughput alternative to traditional animal models before clinical trials ^[23]^. Most PDO-based studies so far report relatively limited sample sizes from single-timepoint sampling (at biopsy or resection) ^[24–26]^.

Organoid culture is more cost-effective than traditional PDX models ^[27]^. To reduce the cost of organoid cultivation and drug screening, research groups even have recently implemented artificial intelligence algorithms to assess organoid vitality and identify specific phenotypic parameters ^[28]^. Moreover, organoid technology has significant potential to become an integral part of routine healthcare, as its establishment (1-2 weeks), overall propagation, along with the time required for drug screening, can be completed within a maximum of five weeks ^[19]^. A comprehensive collection of high-fidelity models is necessary to represent the existing spectrum of disease heterogeneity. Such a resource will not only enable the identification and validation of biomarkers but also facilitate high-throughput functional drug screening for treatment discovery and selection.

Traditional biobanking generally comprises collection of snap-frozen or O.C.T. (Optimal Cutting Temperature™)-embedded frozen tissue samples, along with serum, plasma, and/or buffy coats, to support genomic, transcriptomic, and proteomic research. We established such a biobank within the Division of Thoracic and Upper Gastrointestinal Surgery (TUGI) at the Montreal General Hospital (MGH) of the McGill University Health Centre (MUHC) in 2007 and have amassed a large amount of high-quality non-viable material (single-timepoint and longitudinal) from malignant and case-matched disease-free sites of >2,000 consenting patients, together with comprehensive clinical annotation and follow-up. This resource supported identification of sub-cohorts based on tumor subtypes and other clinical features, building of tissue microarrays, extraction of high-quality nucleic acid libraries for genomics and transcriptomics, and characterization of other biospecimens. To enable translational studies, we expanded our biobanking approach in 2018 to include the simultaneous generation of matched living patient-derived models (PDMs) as extensive tools for target validation, biomarker discovery, and assessment of treatment response (**Supplementary Table 1**).

## 2. Results and Discussion

### 2.1. Patient-derived model (PDM) generation

Between January 2018 and December 2023, we employed live-banking approaches as outlined in **Figure. 1A** and described in the SOPs (Supporting information) to generate 358 PDMs (227 PDOs, 131 PDXs, 5 PDXOs) from 382 patients diagnosed with gastroesophageal adenocarcinoma (**Table 1**). A total of 553 2D-stromal cell cultures were also established from these patient tissues.

**Figure 1.**
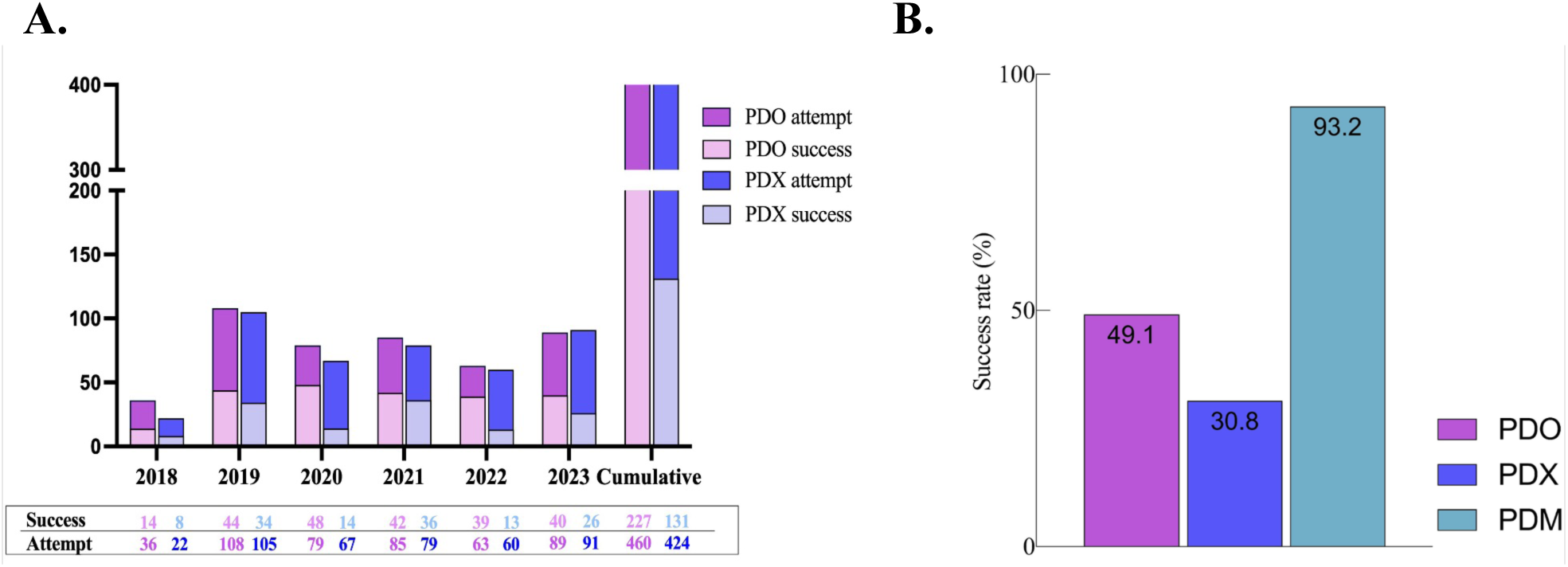
Overview of large-scale GEA Live Bank. We achieved over 90% success in capturing patient-derived cellular models (PDMs) A. Bar graphs showcase annual figures (between 2018 and 2023) for successfully generated PDMs. Numbers of success and attempts are provided. B. The bar graph displays the overall success rate of PDO, PDX, and PDM by sample. This exceeds the success rate of either PDO or PDX alone due to the combined advantage of the dual-pronged approach and longitudinal sampling.

**Table 1.** Summary of PDMs generated from patients classified by epidemiological, clinical and pathological categories. MMR, mismatch repair.

| Patient Characteristics | Total GEA patients (n=382) |
| --- | --- |
| Age (median, yrs) | 67 (18-88) |
| Sex |  |
| Male | 291 (76%) |
| Female | 91 (24%) |
| Tumor grade |  |
| Well | 17 (4%) |
| Moderately | 93 (24%) |
| Poorly | 140 (37%) |
| Not available | 132 (35%) |
| Pathologic Stage |  |
| T0-1 | 34 (9%) |
| T2 | 36 (9%) |
| T3 | 120 (32%) |
| T4 | 54 (14%) |
| Not available | 138 (36%) |
| MSS Status |  |
| MSS | 216 (57%) |
| MSI-H | 31 (8%) |
| Not available | 135 (35%) |
| PDO generation (n=227) |  |
| PDO derived from biopsy | 86 |
| **PDO derived from surgery** | **141** |
| **PDX generation (n=131)** |  |
| **PDX derived from biopsy** | **56** |
| **PDX derived from surgery** | **75** |

For 83 patients, longitudinal tumor samples (i.e., treatment-naïve sample at time of initial endoscopic biopsy, as well as matched generally post-neoadjuvant material from subsequent surgical resection) were collected, the remaining had tumor samples collected solely at diagnosis by endoscopy or at surgical resection. The size of patient-derived tissue samples available for biobanking varies greatly; however, the amount of material obtained from resections is generally greater than that obtained at biopsy. Our success rates in generating PDOs have shown stable improvement over time as sample collection and culture protocols were optimized. The success of PDO generation increased from 39% in 2018 to 45% in 2023, peaking at 62% in 2022, reflecting substantial progress over time . This improvement can be attributed to a series of technical refinements. To increase tissue quality, we established a laboratory adjacent to the endoscopy suite and operating room, enabling rapid tissue processing and minimizing ischemic time. In parallel, tissue collection practices were optimized by preferentially sampling from the leading edge of the tumor and avoiding necrotic regions. Ongoing efforts to standardize experimental protocols for GEA tissue processing, optimize media formulations, and refine the composition of transport media have further helped preserve cell viability over extended periods. Together, these adjustments contributed to the increased success rates observed over time.

Detailed protocols are provided in the Supporting Information. Meanwhile, PDXs have shown a more consistent trend, reaching success of 46% in 2021 and remaining above 20% in other years **(Figure. 1A**). The success rate for PDX generation was lower than that observed for PDO, at 31% and 49% respectively (**Figure. 1B**). By attempting both PDO and PDX simultaneously from a given sample, we increase the likelihood of successfully live bank a tumor by 18% for biopsy samples and 30% for resection samples, demonstrating the value on this dual-pronged approach. When considering the combined success of generating either a PDO or PDX from a specific tumor collected longitudinally over the treatment trajectory, the overall chance of retaining at least one patient-derived model (PDM) from a given tumor surpassed 90%. These success rates exceed those reported in previously published GEA organoid studies, underscoring the robustness of our methodology and further emphasizing the value of our comprehensive approach.

### 2.2. Primary tumor histologic and genomic features are conserved in PDMs

Comparison of H&E-stained sections of FFPE material from PDMs and primary tumor tissues (**Figure. 2A**) reveals that cellular and cytological features of the original tumors were preserved in the PDMs (PDOs, PDXs and PDXOs). Importantly, we observed differences in cellular morphology between PDOs derived from grossly normal and GEA tissue samples (**Figure. 2A** left, upper two panels (tumor) vs. lower panel (normal)). While they both could display a cystic phenotype, tumor organoid lines differed greatly in their thickened cell membrane, disorganized structures, and aberrantly shaped lumen as observed by bright-field microscopy. We also assessed whether clinically relevant features of GEA histological subtypes are conserved in PDOs. Microsatellite Instability-high (MSI-H) tumors are characterized by the absence of at least one of four DNA mismatch repair proteins (MLH1, MSH2, MSH6, and/or PMS2). Immunohistochemical staining of PDOs derived from MSS and MSI-H GEA cases revealed that this feature remained consistent between primary tumor and PDO avatar (**Figure. 2B**). We also observed that individual tumor PDO lines can exhibit varied morphologies. For example, we saw solid and dense organoids of different sizes, as well as more cystic and hollow organoids (**Figure. 2B**).

**Figure 2.**
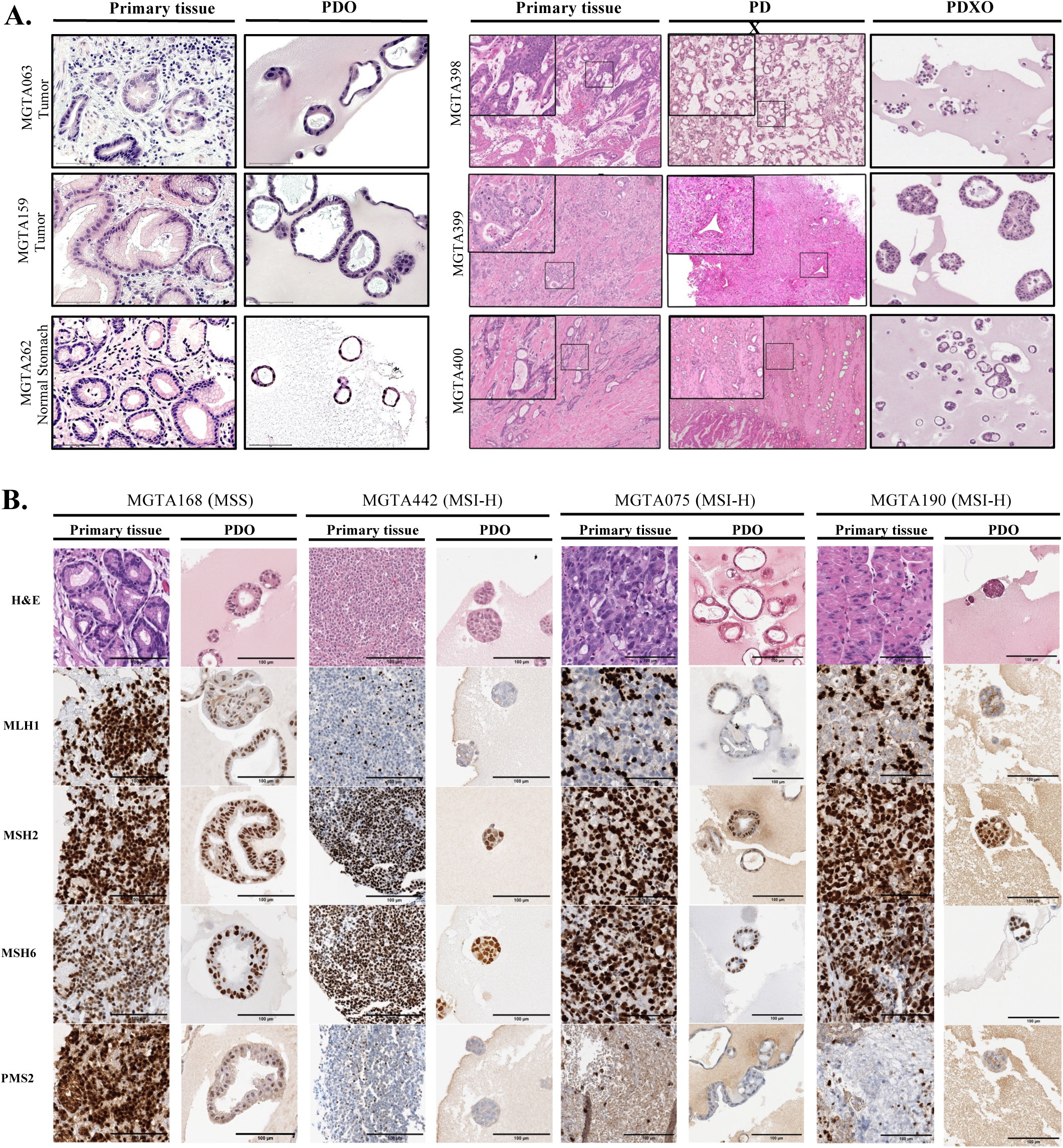
PDOs and PDXs recapitulate histological features of matched primary tissues. A. PDOs (left) and PDXs/PDXOs (right) recapitulate histological features of their tissues of origin (For PDOs: top, resected tumor; middle, biopsied tumor; bottom, resected normal stomach). Scale bar (left panel) = 75 μm. B. GEA microsatellite instability (MSI-H) status is maintained in PDOs. MMR (mismatch repair) protein expression is conserved between primary tissues and matched PDOs – note lack of MLH1 and PMS2 expression in MSI-H case MGTA442 (right). Scale bar = 100 μm.

To evaluate the fidelity of PDO to retain the genomic characteristics of the primary tumor, paired sequencing data from a selected subset of matched primary tumor and organoid samples were analyzed. Concordant driver mutations present in both the tumors and their matched organoids were identified (**Figure. 3**). When focusing exclusively on nucleotide-level mutations, the top recurrent genes include well-characterized drivers of GEA, such as TP53, CDKN2A, and ARID1A. We also compared the distribution of variant allele frequencies (VAFs) between tumors and organoids. The VAFs of concordant mutations were generally retained or increased in PDOs, likely reflecting the higher tumor purity of the cultured models (**Figure. S1**). Notably, driver mutations were predominantly detected at higher allele frequencies in PDOs compared to their corresponding primary tumors. Violin plots stratified by concordance category showed that concordant variants exhibited higher allele frequencies (0.3 – 1.0, average: 0.55), while discordant variants were generally restricted to lower VAFs (0.01 – 0.3, average: 0.15).

**Figure 3.**
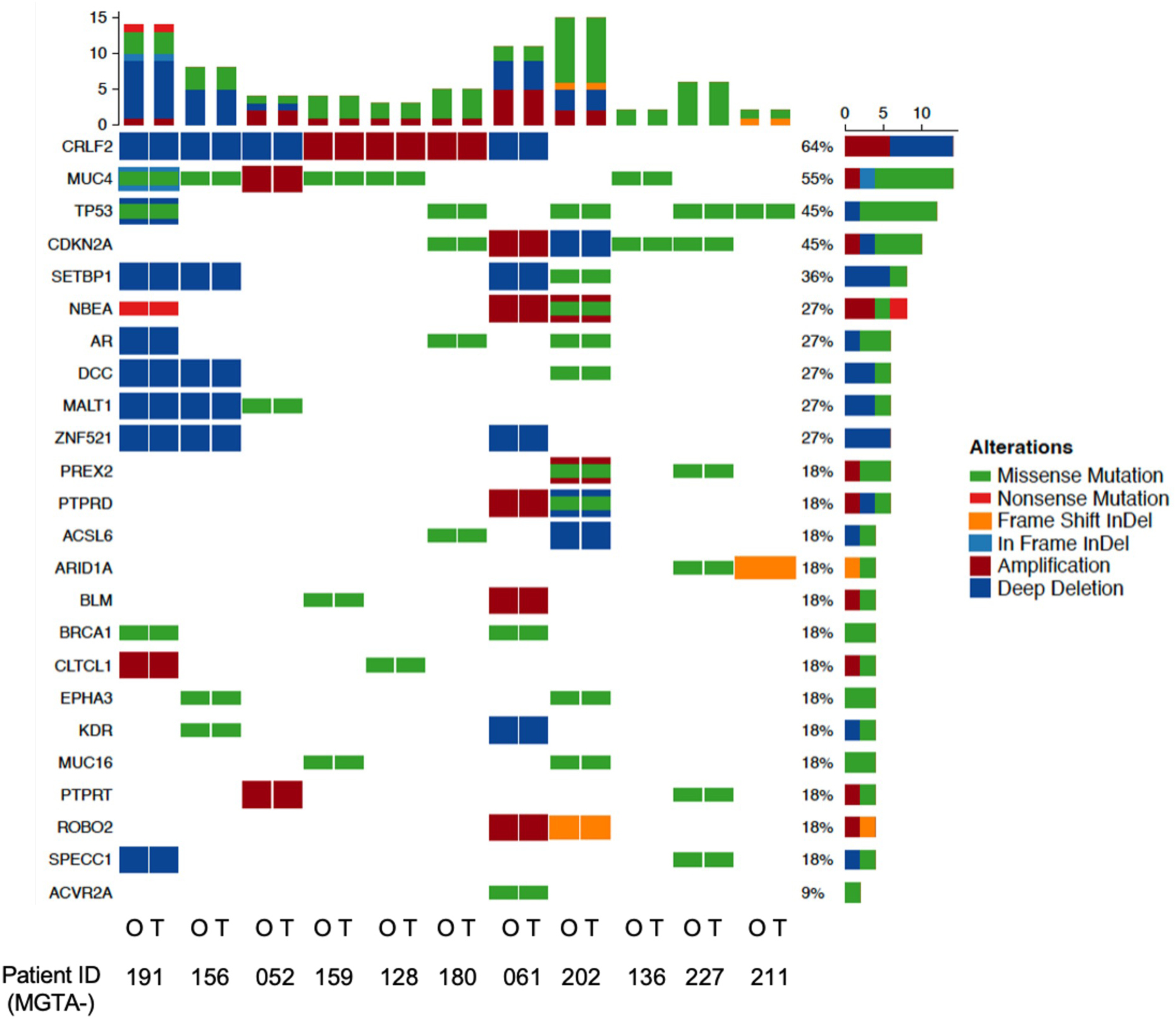
PDOs recapitulate genetic and clonal features of matched primary tissues. An oncoplot of the top 24 mutated cancer genes found in a representative cohort selected from our biobank (11 tumor-organoid paired samples, labeled with tumor as “T” and organoid as “O”).

### 2.3. Determinants of success in PDM generation

We then investigated parameters that may predict the successful generation of a PDO from a given sample. No significant association with tumor stage was observed (**Figure. S4A**); while no association was seen for histological grade for PDOs from biopsy samples, a weak but significant correlation was observed for PDO generation from resection tissues (61.2% for poorly differentiated vs. 48.8% for well- or moderately-differentiated cases; p < 0.05; **Figure. S4B**).

Importantly, no correlation of PDO success with response to NACT was observed (**Table S1**), suggesting that this primarily constitutes a random event rather than being strongly associated with pathological or clinical variables. This is further supported by the observation that successful generation of a PDO from a biopsy did not predict success for PDO generation from the patient-matched resection sample, and *vice versa* (**Figure. S4.D**).

For PDX generation, we observed that success was independent of disease stage; however, histological grade was significantly associated with PDX growth in biopsy but not resection samples (35.8% in poorly or moderately differentiated samples vs. 7.1% in well-differentiated samples; p < 0.005; **Figure. S4.B**). A weak correlation (p < 0.05) with MMR status (across combined biopsy and resection cases) was also observed – this is likely a reflection of the reduced differentiation seen in MSI-H cases. Interestingly, resection (but not biopsy) samples from poor (TRG 3) responders were associated with a higher PDX success rate than those from non-poor (TRG 1 and 2) responders (43.4% vs. 15.9%; p < 0.005; **Table S1**). It is unclear as to whether this represents an example of the link between PDX success and poor outcome (with which NACT response is imperfectly but significantly correlated), or if this simply represents the effect of the increased number of viable epithelial tumor cells expected to be present in samples with poor NACT response.

Sex-based analyses revealed that successful PDO development in biopsy-vs. resection-derived samples are roughly comparable in samples from male patients (42.4% and 54.9%, respectively, **Figure. S4C**), while overall success rates differ significantly in samples from female patients (34.4% for biopsy-derived samples vs 66% for resection-derived samples, p < 0.005).

We also successfully established longitudinal (pre- and post-treatment) PDOs (18 patients) and PDXs (6 patients) (**Figure. S4.Di**). Remarkably, we obtained a total of 46 PDXs (21 from biopsies; 25 from surgical resections) from patients for whom we were unable to establish corresponding PDOs (**Figure. S4.Dii**). Subsequently, we endeavored to create PDXOs from 20 PDXs, achieving success 25% of the time. This finding further supports our decision to proceed with parallel PDX and PDO workflows for each sample when possible.

Subgroup analyses indicate that the combined rate of success is 59% for biopsy samples (vs. 41% for PDO alone and 33% for PDX alone) and 72% for resection samples (vs. 42% for PDO alone and 32% for PDX alone). For our longitudinal approach, in cases where the generation of live preclinical models was attempted at both the biopsy and resection time points, the combined success rate for PDOs per patient was 75% (vs. 40% for biopsy and 46% for resection alone).

Similarly, longitudinal PDX implantation led to a success rate of 64% (vs. 3% for biopsy and 42% for resection alone). Overall, in the cases where both PDO and PDX generation was attempted at both biopsy and resection time points, we achieved at least one tumor avatar in 93% of cases, further justifying our combinatorial approach for situations when a live cancer model is absolutely desired (e.g. rare tumor types) (**Figure 1B**) .

### 2.4. Assessment of PDM purity

The growth of isolated cells as PDOs introduces the possibility of contamination with non-epithelial cells from the original sample. To assess this, we performed fluorescence-based flow cytometry analyses directed against the epithelial cell lineage marker EpCAM (epithelial cellular adhesion molecule). Nearly 82% of cells in the tumor-derived PDO model expressed this marker (**Figure. S3.A**). As PDXOs might be contaminated with murine cells, we also tested these samples. Using flow cytometry with human-specific marker (EpCAM) and murine-specific (H2Kd) markers, it was determined that 77% of PDXO cells were of human origin (**Figure. S3.B**) at early passage.

### 2.5. Isolation and expansion of stromal elements

We additionally isolated, cultured, and banked stromal cell fractions (cancer-associated fibroblasts/CAFs, n=272; tumor-infiltrating lymphocytes/TILs, n=281) from tissue samples as valuable tools to investigate the tumor microenvironment (TME). To evaluate the purity of the isolated CAFs, we conducted immunofluorescence (IF) staining directed against the fibroblast markers aSMA and/or PDGFRβ (**Figure. S3Ci**). Furthermore, we utilized flow cytometry to quantify cells expressing the fibroblast markers FAP, CD74, PDGFRβ, and PDGFRa in our CAF cultures (**Figure. S3.C.ii**) to confirm the overall purity of the preparation. Similarly, we employed flow cytometry targeting the immune cell lineage marker CD45 to ensure that the majority of cells cultured as TILs maintained their immune lineage integrity even after 3-4 passages (**Figure. S3.Di-ii**). Additionally, we established two co-culture models using PDOs combined with these stromal cell fractions (**Figure. S3.Ciii, S3.Diii**). These high-fidelity models constitute valuable resources for the further study of stroma-driven resistance mechanisms.

### 2.6 Treatment of PDOs *in vitro* mirror in-patient tumor response to chemotherapy

Key downstream applications for PDMs in the field of precision oncology include prediction of intrinsic response to various therapies (e.g. chemotherapy, targeted therapy, radiation, immunotherapy) and testing of alternative therapies within a clinically useful timeframe. We collected tissues at specified time points during neoadjuvant therapy (**Figure. 4A**). We assessed whether established PDOs recapitulate the treatment pathological response observed in their “patients of origin” at the time of resection, as defined by pathological Tumor Regression Grade (TRG; 0, complete; 1, near complete; 2, partial; 3, absent). In a selected cohort of 15 PDO lines derived from patient tumors prior to systemic treatment, including both “good” (TRG0/1) and “poor” (TRG2/3) responders to subsequent docetaxel-based triplet chemotherapy, were subjected to the same regimen as that received by the patients in vitro for 72 hours (**Figure. 4B**). Each PDO line was tested in six replicates, and dose-response curves were then generated based on post-treatment viability measurements. Notably, PDOs derived from chemotherapy-resistant patient tumors exhibited significantly higher viability across multiple concentrations (p<0.0001) (**Figure. 4C-D**).

**Figure 4.**
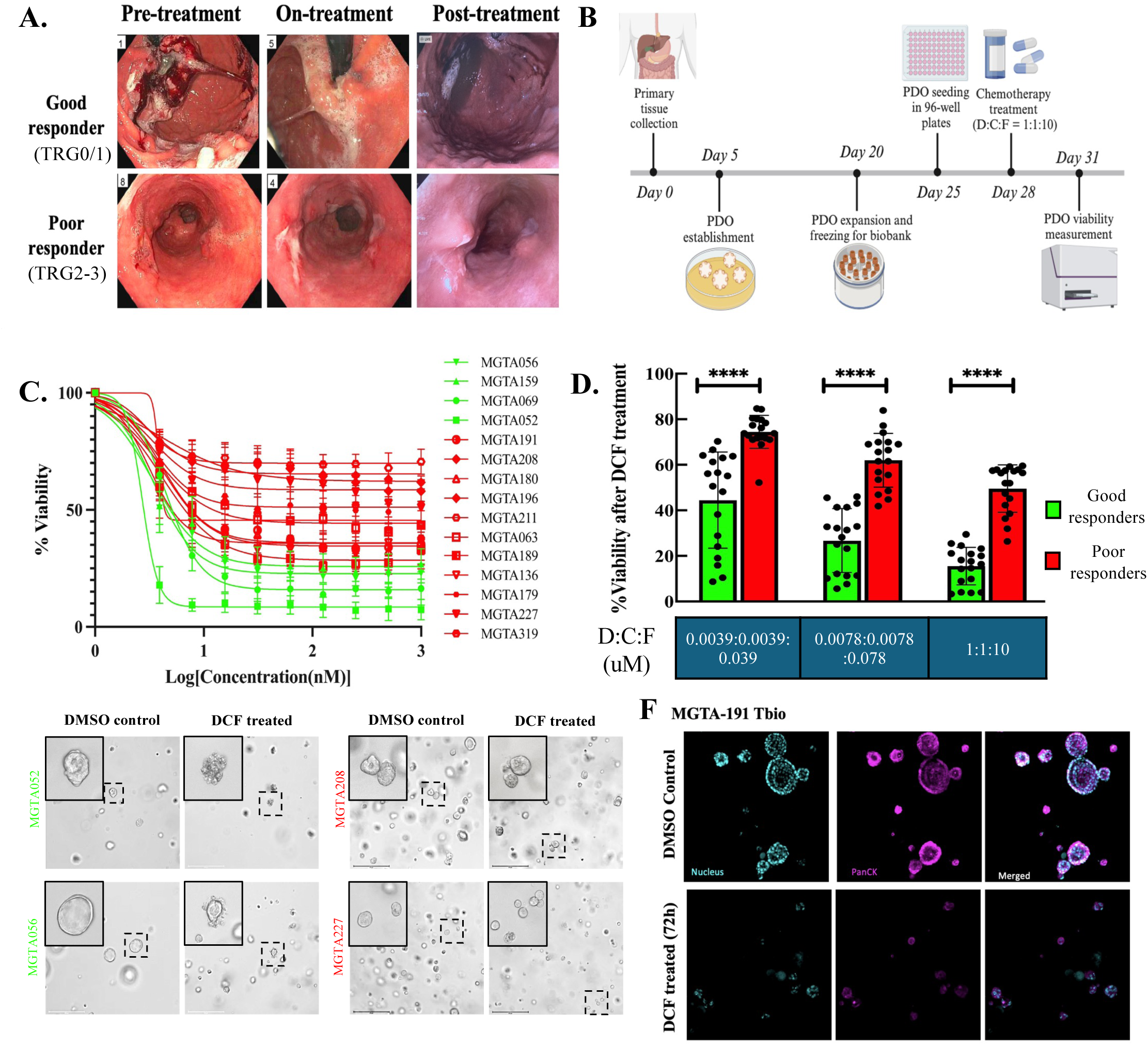
PDOs derived from treatment-naïve biopsies recapitulate in-patient chemotherapy response upfront. A. Endoscopic images captured at three longitudinal timepoints for GEA patients who received neoadjuvant chemotherapy as SOC. Patients categorized as good responders exhibit TRG 0/1, while those classified as poor responders have TRG2/3. B. Timeline of PDO generation and in vitro evaluation of efficacy of DCF treatment. C. DCF response curves of PDOs derived from 4 good responders and 11 poor responders. Red: poor responders PDOs; Green: Good responder PDOs. D. Bar graph showing that the viability of PDOs from good responders is significantly lower than that of PDOs from poor responders after DCF treatment at three critical concentrations. E. Brightfield images depict the morphological alteration of PDOs following DCF treatment. F. Immunofluorescence images depict epithelial marker PanCK expression changes in PDOs following DCF treatment. Multiple t-test is used to compare the cell viability between two groups in panel D, ****p<0.0001.

Organoid morphology was assessed using brightfield imaging. Consistent with higher post-treatment viability, PDOs derived from chemotherapy-resistant tumors exhibited relatively intact cellular architecture. In contrast, PDOs from patients classified as good responders showed marked morphological changes and reduced PanCK expression, indicative of apoptosis and membrane rupture (**Figure. 4E–F**). This highlights the effectiveness of endoscopy-derived chemo-naïve PDOs in predicting chemotherapy response.

Incorporating longitudinal sampling and PDO generations into our biobanking pipeline enables us to examine the treatment response trajectory over time and live models of both innate and acquired resistance. We collected primary tissue from the same patients at the time of initial biopsy and after neoadjuvant chemotherapy (at resection) and generated matched PDOs. Biopsy-derived organoids from patient MGTA-159, classified as a subsequent good responder (TRG 1), demonstrated sensitivity to DCF treatment *in vitro* (IC_50_ = D: 3.37 nM, C: 3.37 nM , F: 33.7 nM; normalized AUC = 0.39). However, when their post chemotherapy-primed, surgical resection-derived PDOs were exposed to DCF *in vitro*, acquired resistance was observed, as indicated by a higher IC_50_ of D: 5.85 nM, C: 5.85 nM, F: 58.5 nM; and normalized AUC of 0.51 (**Figure 5.A**). A similar pattern of heightened resistance to chemotherapy was observed in longitudinal PDOs derived from poor responder (TRG 3) MGTA-180. In this case, the biopsy-derived PDOs exhibited intrinsic resistance. In addition, the matched resection-derived PDOs displayed further elevations in IC_50_ and AUC compared to their biopsy-derived PDOs (IC_50_: D: 4.11 nM, C: 4.11 nM, F: 41.1 nM *vs.* D: 5.83 nM, C: 5.83 nM, F: 58.3 nM, normalized AUC: 0.48 vs. 0.53) (**Figure 5.B**). This pattern of chemoresistance (innate/intrinsic and acquired) was consistent with the clinical trajectory of these patients, as both experienced cancer recurrence within 8 months of surgery (**Figure. 5**). Consistently, comparison of four additional biopsy- and resection-derived PDOs demonstrated higher resistance to docetaxel-based triplet therapy in the resection-derived models, as reflected by increased normalized AUC values (**Figure. S3**), supporting the emergence of acquired chemoresistance following neoadjuvant treatment.

**Figure 5.**
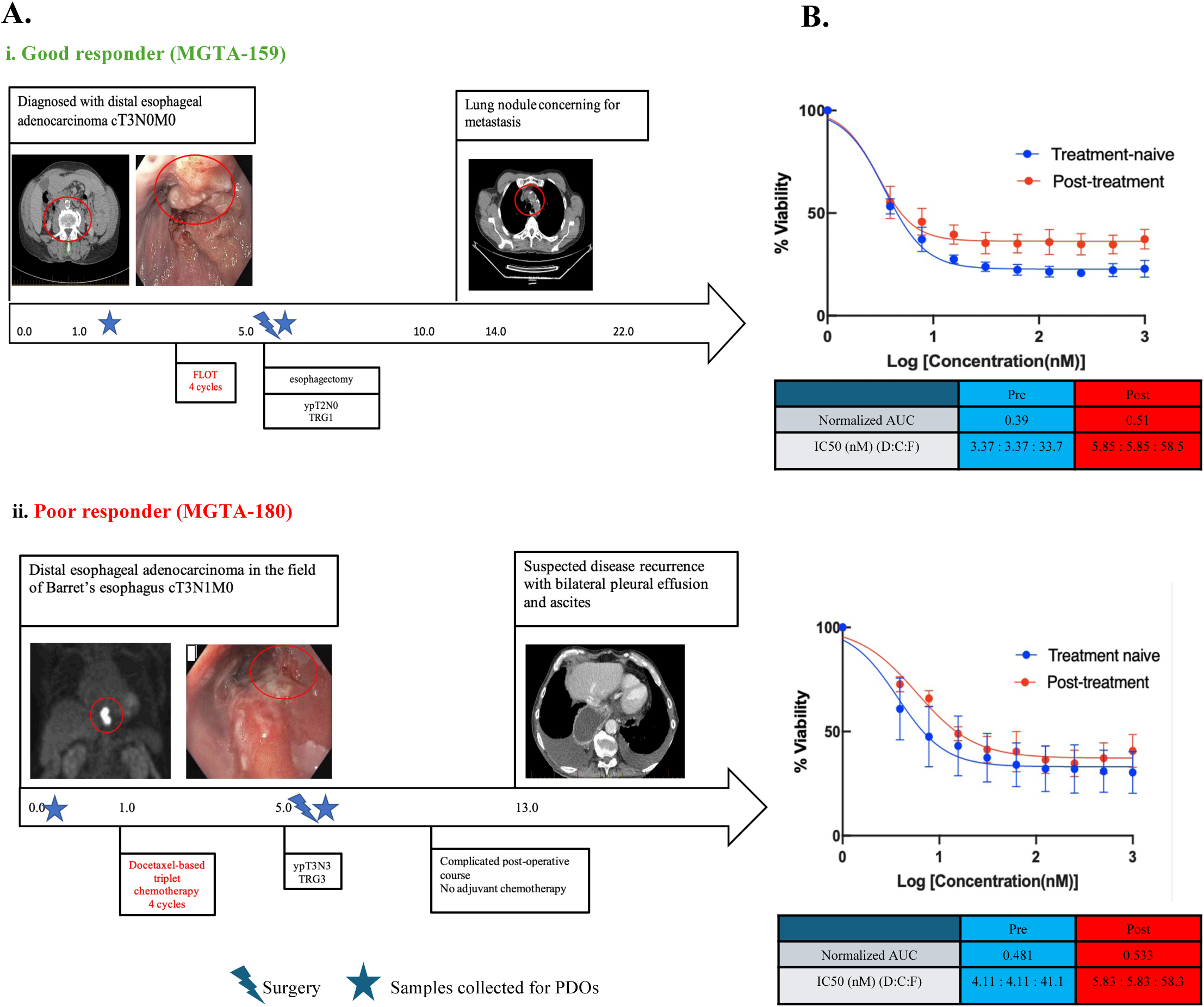
PDOs recapitulate longitudinal treatment response. A. i-ii) Treatment timeline of a chemo-sensitive and chemo-resistant patient. The numbers within arrows represent months since initial diagnosis. A star marks the timepoints at which samples were taken for PDO generation, while a thunderbolt symbol indicates surgical resections. B. Survival curves against DCF treatment using PDOs from patients categorized as good- and poor-responders. The PDOs were derived from same patient tissues collected at two different stages: pre-treatment (diagnostic biopsy) or post-treatment (surgical resection). PDOs derived from post-treatment tissues exhibit higher resistance levels, characterized by a greater Area Under the Curve (AUC) and IC_50_ values (GraphPad), highlighting increased chemo-resistance. GraphPad), highlighting increased chemo-resistance.

### 2.7. Generation of a chemotherapy response profile of a subset of GEA PDOs from chemo-resistant tumor

To demonstrate the utility of GEA PDOs as a foundation for a functional precision oncology platform, we sought to develop a chemogram to identify potential salvage therapeutic approaches for chemo-resistant tumors. High-throughput single-drug screening was performed for five PDOs derived from chemo-resistant patients with a panel of 22 selected drugs from a library of 3198 FDA approved/in-trial drugs (**Figure. 6**). The chemogram results indicate that each patient-specific PDO shows a unique drug sensitivity pattern across 22 selected chemotherapy drugs. The mutational profiles of representative oncogenes demonstrate that there was no clear association between drug response and genetic alteration pattern. Despite this genetic heterogeneity, we observed that idarubicin, vincristine, and gemcitabine were the three most common efficacious chemotherapy drugs other than docetaxel that caused <50% cell viability across all five PDO lines. In the chemogram, the median number of hits (<50% cell viability) per PDO was 13 (range: 8-19). Among the 22 drugs, in addition to the SOC for GEA (docetaxel, oxaliplatin and 5-fluorouracil), 19 drugs were identified as candidate hit drugs in at least one PDO line, and they were FDA approved for treatment of various types of cancers or undergoing clinical trials for GEA patients (**Supplementary Table 4**). This suggests that a chemotherapy drug originally developed for non-GEA cancers could serve as a patient-specific treatment option for chemo-resistant GEA tumors, opening unexpected and valuable drug-repurposing opportunities.

**Figure 6.**
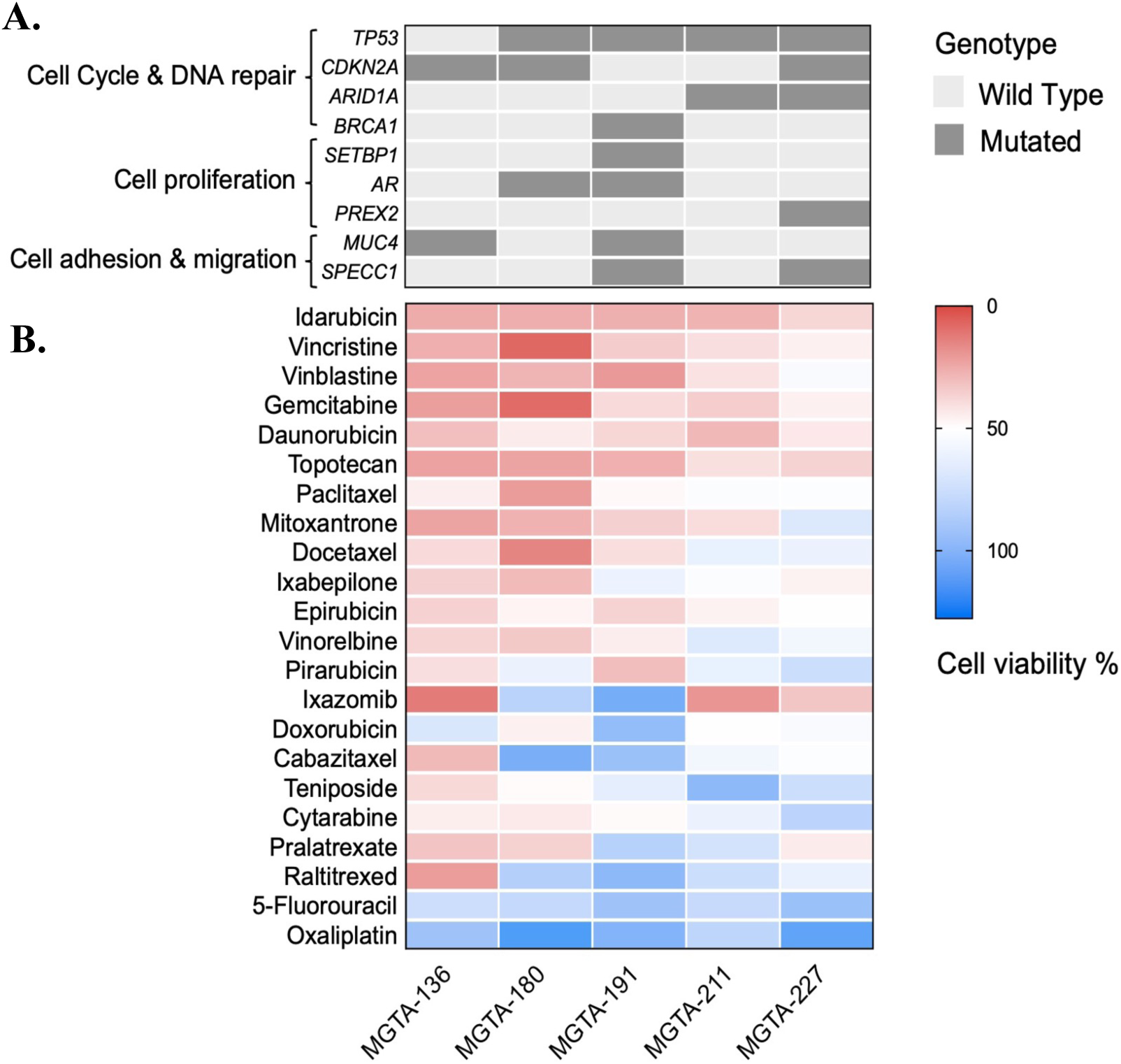
Identification of alternative chemotherapy options in the context of heterogeneous genomic alterations. A. Representative genomic mutation profiles and their associated pathways of five PDOs derived from chemo-resistant patients. B. The heatmap presents the mutational status of selected genes in conjunction with a chemogram, a widely used approach for guiding the selection of chemotherapy agents for individual patients. Representative chemogram indicates the personalized responses (expressed as percentage of cell viability) to 22 FDA-approved or investigational chemotherapy drugs across five PDO lines. PDOs were exposed to 1 μM of each drug for 72 hours, and cell viability was determined using the CellTiter-Glo 3D assay.

## 3. Conclusions

GEA, with an increasing mortality rate worldwide, continues to be a significant public health concern. Most patients experience a poor outcome (< 20% overall survival at 5 years).

Terminally differentiated 2D cell line models display reduced fidelity due to a lack of heterogeneity. Simultaneously, spontaneously arising, genetically modified animal models are absent due to substantial structural and morphological differences between the human and murine esophagus. Clinically relevant models that can be used for basic research, drug/biomarker discovery and real-time clinical decision-making in GEA are crucial to improving personalized treatment and patient survival in this poor-outcome disease, where only incremental advances in therapy have been observed in the last 25 years.

To this end, we have developed a comprehensive large-scale biobanking approach to incorporate PDMs as tumor avatars, enabling downstream investigations of tumor heterogeneity, clonal expansion, behavior and sensitivity to both standard-of-care and investigational agents. Leveraging previously published methods and iteratively optimizing them, we developed novel culture protocols for generating both 2D and 3D models from tumor (n=227) and tumor-adjacent normal (n=203) tissue-derived PDOs using a modified, growth factor-reduced Matrigel-based 3D cell culture system ^[19, 29]^. A total of 272 matched CAF and 281 TIL cultures were also generated. This unique resource facilitates stroma-inclusive co-culture models that may better model tumor response to therapy, as we have recently demonstrated with a patient-specific esophageal adenocarcinoma organ-on-a-chip ^[30]^. Based on histopathological, genomic and drug-response parameters, these PDO avatars closely recapitulate key features of their parental tumors.

Our live-banking and tumor model generation is closely integrated with collection of relevant clinical information. Combining clinical information and large numbers of tumor avatars permits focused investigations into sub-cohorts of patients that are only possible in large-scale initiatives including: various histological subtypes of GEA such as MSI-H; sex-dependent variability; tumors of different stages and grades; and differential response to systemic therapy (e.g. chemo-sensitive vs chemo-resistant). This latter distinction represents a key element in our ongoing initiative to discover biomarkers predictive of response, as well as vulnerabilities in non-responder samples that can be targeted to improve treatment efficacy.

Current 3D disease model-based research often encounters limitations such as small sample sizes and low success rates. Crucially, our ability to achieve consistently high success rates in live tumor banking across hundreds of GEA patients reflects a series of integrated and sustained technical improvements. In the area of tissue collection, we established a laboratory adjacent to the operating sites, and refined sampling protocols that prioritize viable, non-necrotic regions.

We also optimized media formulations in-house, adjusting concentrations of individual components and incorporating additives such as N2 supplements to support the viability and long-term expansion of GEA-derived cells. In parallel, standardized SOPs were developed and continuously refined to ensure reproducibility over the years. Once these protocols were in place, we were able to extend our workflow to accommodate tissue collected remotely, supported by the use of our optimized transport media. This capability now enables us to receive samples from external institutions, expanding the reach of our biobanking effort. To our knowledge, this represents the largest living biobank for GEA to date, offering a valuable resource to the broader research and clinical community.

In addition to progressive technical refinement, we implemented a dual-model (PDO/PDX), and dual timescale (longitudinal: pre and post treatment) workflow to substantially enhance success rates in generating personalized tumor avatars for patients with GEA culminating in a >90% success rate in generating at least one PDM for an individual patient. This is significantly higher than what has been reported for single-approach studies. Published GEA PDO studies have reported establishment rates ranging from approximately 31% to 78%, with an average success rate around 65% ^[24–25, 31–33]^. While some of these success rates are comparable to ours on a per-tumor basis, most prior efforts have been constrained by small sample sizes, typically generating between 9 and 73 PDOs in total. In contrast, our biobank has successfully established over 400 PDO lines as part of the effort described here. PDX engraftment rates tend to be lower across gastric and esophageal cancers; a recent single-timepoint PDX study reported the generation of 100 gastric adenocarcinoma models from 349 fresh surgical samples (29% success rate). Taken together, these comparisons underscore the robustness of our optimized live-banking strategy - not only in achieving a high overall success rate, but also in sustaining this performance at scale. The formation of PDMs appears to be primarily a random event - the successful generation of a PDO line from a sample is not correlated with growth of a matched PDX sample, and vice versa, or with most of the clinical variables investigated here. Similarly, success in PDO generation from a patient sample taken at one point in their clinical time-course does not predict success for a sample taken at another timepoint. The factors underlying these differences are unclear – one potential explanation is that since the samples for PDO vs. PDX are physically distinct, as are those taken at different timepoints, only a subset of the tumor, as defined by physical location, is capable of PDM formation. Further studies involving parallel processing of multiple samples taken from different areas of the same tumor will be required to answer this question. The observation that PDX but not PDO generation is correlated with NACT response is of interest; success for good vs. responders (TRG 1 and 2 vs TRG 3) is significantly lower in PDXs, while PDO generation is independent of this variable. This further accentuates the importance of our dual approach, which allows for the capture of a more representative cohort encompassing the full spectrum of patient response as PDOs.

Importantly, our approach enables screening of a range of potential therapies, including SOC chemotherapy, within a 31-day timeframe. Since the standard 4 cycles of neoadjuvant therapy typically span 10-12 weeks, this allows for the evaluation of treatment response and the potential incorporation of validated supplementary agents within a clinically relevant window.

Additionally, when evaluating treatment responses in longitudinally derived PDOs from the same patients, our findings reveal that despite originating from the same patients, these PDOs exhibit diverse responses to DCF treatment, mirroring their clinical context and downstream outcomes. The longitudinal generation of PDOs and comparative analysis of their drug responses not only facilitates studies of how treatment response evolves over time and the discovery of mechanisms underlying chemoresistance but also enables independent predictions for both neoadjuvant and adjuvant chemotherapy response. In addition, simultaneous isolation and culture of stromal elements including TILs and fibroblasts permits assembly of models incorporating the contributions of the tumor microenvironment, which is critical for the evaluation of new modalities that target this combinatorial milieu (e.g., immuno-oncology). The large size of the model cohort generated here permits investigation of the efficacy of large drug panels across hundreds of samples, facilitating the identification of agents efficacious in small subpopulations. In combination with extensive profiling of these samples at the transcriptomic, genomic and proteomic levels, biomarkers predictive of response to specific novel agents (either alone or together with SOC) can be discovered and these biomarker/drug combinations can be validated in additional samples, finally bringing the promise of personalized medicine to the field of GEA. In summary, we present a targeted, scalable workflow for the collection and maintenance of both viable and non-viable GEA samples. These PDMs, which on a per-patient basis can be generated for nearly 95% of cases, effectively recapitulate key histological, genomic, and drug-response features of their parental tumors, making them invaluable tools for conducting in vitro functional assays and assessing responses to various treatment modalities, including SOC and alternative therapies.

## 4. Experimental Section

### Sample selection

Generation of PDMs is highly resource- and labor-intensive. Due to limited resources, it is not feasible for our group to live-bank samples from every GEA case that undergoes biopsy or resection at the MUHC. We have defined a set of criteria for PDM stream selection based on our group’s primary research focus. These include grossly or histologically diagnosed GEA lesions located within the digestive tract from the distal esophagus to the proximal stomach. Patients who received CRT (chemoradiation therapy) are excluded since this treatment approach is rare for GEA at our center. As a tertiary treatment center, the TUGI service of the MUHC often performs surgical resections for patients whose initial biopsy and diagnosis of GEA occurred at other sites. Since collecting longitudinal samples across the treatment continuum creates added value, we prioritize cases whose initial diagnosis was performed at the MUHC for live banking when possible. Absolute priority for tissue allocation must be given to clinical pathology needs, and thus fresh tissue is not always available for biobanking for smaller lesions. Notably, patients showing excellent response to SOC therapy may have no residual tumor at the primary site post-treatment, necessitating the omission of such samples from research collections.

### Sample collection

An integrated team of clinical and research personnel identifies appropriate candidates, obtains patient consent before sample collection, and longitudinally (pre-/on-/post-treatment) collects fresh tissues as well as additional biospecimens (blood, ascites, etc) from biopsies/surgical resections. Tissue and blood sample collection is closely associated with clinical care to minimize the burden on patients. At biopsy, several additional samples are taken by the endoscopist after all pieces required for clinical diagnostic purposes have been collected. These additional samples collected specifically for live biobanking efforts are taken preferentially from the leading edge of the tumor avoiding any necrosis and immediately placed in tubes containing transport medium (see Supporting Information) and transferred at 4°C to the processing site. At surgery, resected tissues are immediately evaluated by the on-duty pathologist in the surgical pathology suite. If there is excess material beyond clinical requirements, small pieces identified as normal and/or tumor tissue by the pathologist, along with potentially metaplastic or dysplastic regions when available, are taken for biobanking. These pieces are then placed in ice-cold transport medium for subsequent processing, typically within 15-20 minutes. Importantly, close integration and short transport distances (same building; 2-floor distance for resections, 3-floor for biopsies) minimize warm ischemia times. However, we have more recently employed this approach and media to successfully generate organoids at a high success rate (14/21 = 67%) from tissue collected remotely (e.g., >2000 km distance) and transported overnight to our center for processing, as described below. Notably, among these, biopsy samples achieved a success rate of 80%, highlighting the robustness of our protocol even under suboptimal transport conditions (data not shown).

### Sample processing and cell culture

At the processing lab, tissue is divided into multiple pieces based on project requirements. Freshly flash-frozen, O.C.T.-embedded, and FBS/DMSO (viable frozen) fragments of each tissue type are generated and stored at -80°C or in liquid N_2_ as appropriate. For live-banking sample allocation, project-specific considerations are the primary criteria, with a priority given to generating PDOs/TILs/CAFs over PDXs. For PDX (and subsequent PDXO) generation, 1-2 (depending on sample size) small tissue pieces are implanted subcutaneously in NOD/SCID nude mice (Strain #005557). The remaining tissue is mechanically and enzymatically digested to single cells and used to generate PDOs and 2D cell models (TILs and fibroblasts).

A significant part fresh tissue was minced and enzymatically digested with tissue digestion buffer in a GentleMACSTM Octo Dissociator with heaters (Milltenyi Biotec.). Very small pieces from tumor regions were used separately for TIL culture in expansion media (StemCell Cat No. 10981) supplemented with human recombinant IL-2 (CHO-expressed) (final conc. 10^3^ CU/mL) and ImmunoCult™ Human CD3/CD28/CD2 T Cell activator (StemCell Cat No. 10970). The cell pellets were collected and trypsinized to obtain single cells. The resulting cells were resuspended and were let stand for 10 min. Later a fraction of the upper part of the supernatant was transferred to a new tube for fibroblast culture. PDOs, TILs and associated fibroblasts were all cultured under hypoxic conditions, i.e., 3% O2. For organoid culture, cell pellets were resuspended in ice-cold Matrigel (Corning 356255) and seeded on 24-well plates. The GEA organoid medium was modified using published protocols and commercially available components. The composition and concentration of all medium components were optimized in-house to support the efficient establishment and long-term expansion of GEA-derived PDOs.

Specifically, we replaced the original Advanced DMEM/F12 supplemented with 10% FBS with the commercially available IntestiCult™ Organoid Growth Medium (STEMCELL Technologies, #06010), which provides a more standardized, serum-free environment. To further support epithelial lineage maintenance and culture stability, we added N2 supplement (Thermo Fisher Scientific, #17502-508) to supply additional micronutrients. Cell lines used to produce Wnt3a/R-spondin/Noggin conditioned media were generously provided by Dr. Hans Clevers (Hubrecht Institute, Netherlands), Dr. Calvin Kuo (Stanford University, USA), and Dr. Catherine O’Brien (University of Toronto, Canada). Detailed medium compositions are provided in Supplementary **Tables 2 and 3**. Organoids in Matrigel domes were maintained in expansion media. Mature PDOs were passaged after 10–14 days. Single cells were obtained through enzymatic digestion and trypsinization. For propagation of fibroblasts, cell pellets were resuspended in 1 ml fibroblast propagation media and seeded in Type I collagen precoated plates. Detailed experimental steps and reagents used are described in the Supplementary Information . Detailed experimental steps and reagents used are described in the Supplementary Information.

In selected cases, the largest fraction of the isolated single-cell suspension is immediately subjected to single-cell RNA sequencing (scRNAseq). Data obtained from scRNAseq are not shown here. We also prepare organoid histogel blocks following an existing protocol (https://ccr.cancer.gov/sites/default/files/2022-11/Histogel_Protocol_0.pdf) to facilitate immunohistochemical analysis and supports additional biomarker discovery and validation studies.

### Genomic sequencing and analyses

For genomics assays of resection samples, frozen primary tissue is sectioned, stained and verified by experienced clinical pathologists; tumors with over 30% purity are selected for further processing. DNA is extracted from primary tumor tissue, blood samples and corresponding PDOs using a Qiagen kit (Cat No. 80204) and sent to a dedicated genomics facility (Genome-Quebec) for library preparation and whole exome sequencing (WES) using Illumina NovaSeq technology. WES data were processed using the NextFlow pipeline Sarek to generate analysis-read bam file ^[34]^. Within the pipeline, following BWA alignment ^[35]^of paired-end reads using hg38 as the reference genome, we used GATK 4.2.3.0 ^[36–38]^ to process the raw read fastq files and obtained analysis-ready bam files. The processing steps followed guidelines of GATK best practice ^[39]^. Variants were called using GATK’s Mutect2 program. called variants were subsequently filtered using FilterMutectCalls.

Variant annotation and visualization with oncoplot. Called and filtered variants were saved in .vcf format and subsequently annotated with Annovar ^[38]^, supplying the -protocol option with the following arguments: -protocol refGene, cytoBand, exac03, avsnp147, dbnsfp30a, 1000g2015aug_all, cosmic70. We then used vcf2maf script ^[40]^ provided by the Sloan-Kettering Institute to convert each annotated file into maf format. Multiple maf files were relabeled with “Tumor_Sample_Barcode” and combined using in-house python scripts.

Consensus cancer genes for the validation step were obtained from COSMIC ^[41]^. Variant allele frequencies were calculated counting the number of reads that contain each allele in the VCF file, which was generated by Mutect2. Driver mutations were selected from the mutated consensus cancer genes that have VAF higher than 0.05. In addition, we recorded cancer genes that have copy number variant (CNV) events with log2 values beyond ±0.75, respectively.

Concordant mutations were identified by taking the intersection of the mutations in tumor and organoid. Discordant mutations are the ones that are not in common between tumor and organoid. Using the oncoprint () function in ComplexHeatmap package in R ^[42]^ we visualized the top 24 concordant driver mutations and copy number variants among the 11 tumor-organoid paired patient samples.

### Common cancer gene mutation recapture rate analysis

From the data summary of Maftools, we count the number of mutations among the top 25, 50, 80, as well as all known cancer genes. The similarity metric Jaccard index is used to measure the rate of cancer mutation recapturing:

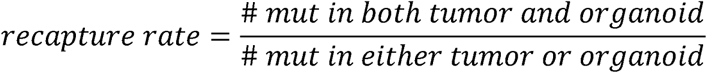

Similarly, we define the consistency rate as the fraction of mutations that identified in organoid and were also consistently found in tumor samples:

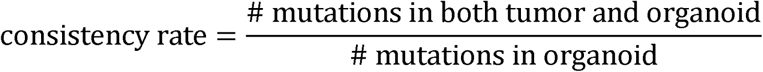

### Pairwise similarity matrix analysis of cancer gene mutations

All 11 paired samples were compared in pairwise fashion with known cancer gene mutations called. For each mutation, when the Hugo symbol, chromosome number, start position, and mutant classification all match, a match is counted for the pair in question. All but the self-pairs were examined for each known cancer mutation called. Similarly, we performed the same analysis with all known SNP called in the sample swap test.

### Tumor mutation burden calculations

We used the tcgaCompare() function from maftools R package to compute the tumor mutation burden in primary tumors, organoids, and the combination of the two sets of samples. All parameters were set following the default settings on the webpage of maftools documentation ^[43]^.

### Clinical correlations

GEA biopsy and resection tissue samples (as formalin-fixed/paraffin-embedded (FFPE) material/hematoxylin and eosin (H&E)-stained sections) are assessed in the clinical pathology setting to derive relevant information including tumor histology, grade, stage, and response to therapy. Immunohistochemistry (IHC) or fluorescence in situ hybridization (FISH)-based assays define parameters such as MSI (microsatellite instability) status, HER2 amplification or histological subtyping (e.g., adenocarcinoma vs, squamous cell carcinoma) - currently, genomic testing is not routinely carried out for GEA at our site. All of this information is captured in our research database, along with clinical information regarding history, symptoms, clinical staging and investigations, treatment, response, and follow-up as well as extensive quality-of-life data. This patient-specific information is coded and entered into a customized digital platform (REDCap) ^[44]^, facilitating annotation and data sharing while minimizing risks to privacy and data loss.

### In vitro drug treatment on organoids

10000 cells were resuspended in 10uL Matrigel on ice and seeded in each well of a 96-well plate. The plate was left in 37°C incubator to solidify the Matrigel, followed by addition of 200uL of complete growth media (1:1 IntestiCult media and Gastric media, see Supporting Information for details) into each well. Organoids were allowed to grow for 3-4 days until they display mature morphology. Docetaxel (Taxotere, Sanofi-Aventis Canada), 5-Fluorouracil (Cayman Chemical Company, #14416) dissolved in DMSO, and cisplatin (Cayman Chemical Company, #13119) dissolved in sodium chloride solution in water were dispensed to each well using a Tecan D300e Digital Dispenser (Tecan) at a 1:1:10 (D:C:F) ratio. 72 hours after treatment, organoid viability was measured using CellTiter-Glo 3D reagent (Promega, G9681) according to the manufacturer’s instructions. Luminescence readings were obtained from a Varioskan LUX Multimode Microplate Reader (ThermoScientific). Readings were normalized to DMSO-alone vehicle control. Viability curves and IC_50_ values were generated using GraphPad software.

### High-throughput drug screening

A library of 3198 compounds from FDA Approved & Pharmacopeial Drug Library was purchased from MedChemExpress to conduct high-throughput drug screening. Furthermore, a panel of 22 chemotherapeutic agents was selected from this drug-library for the preparation of the chemogram. In 15 µl 70% matrigel, 800 cells were seeded on 384-well plates. Cells were cultured for 3 days in 40 µl complete media to allow maturation of organoids, at 37°C in a humidified atmosphere in a 5% CO2/3% O2 tri-gas incubator. Then PDOs were dosed at 1 µM of the compounds for 72 hours. Cell viability was measured using CellTiter Glo 3D reagent according to the manufacturer’s instructions. Quantification of luminescence readings was obtained using a Varioskan LUX Multimode microplate reader. Cell plating, drug treatment, and viability assay were all conducted by an automated liquid handler (Beckman Biomek i5) to ensure accuracy. The detailed clinical information associated with the 22 drugs shown in the chemogram is listed in Supplementary **Table 4**.

### Statistical Analysis

Analyses for *in vitro* assays were conducted using GraphPad Prism version 10.1.1 for MacOS, GraphPad Software (www.graphpad.com). Any *p* values <0.05 were statistically significant.

Statistical significance for biobanking was determined by Chi-squared test or multiple unpaired t-test.

## Supporting Information

Supporting Information is available online or from the author.

## Acknowledgments

We would like to extend our sincere thanks to the participants who kindly provided consent for the use of their samples and data in this project. This work was made possible by generous support from the Jarislowsky Foundation *via* the Montreal General Hospital Foundation, as well as grants from the Cancer Research Society, the Canadian Institutes for Health Research, a Cancer Research UK Grand Challenge and the US Department of Defense/Congressionally Directed Medical Research Programs. We would like to acknowledge technical assistance from the Immunophenotyping, Histopathology, Molecular Imaging and Biobank Technology platforms (Research Institute of the McGill University Health Centre); the Histology platform of the Goodman Cancer Institute and the Advanced BioImaging Facility of the the Life Sciences Platform (McGill University) and the Genomics/Sequencing Platform (Genome-Quebec). We also thank Dr. Hans Clevers (Hubrecht Institute, Netherlands), Dr. Calvin Kuo (Stanford University, USA), and Dr. Catherine O’Brien (University of Toronto, Canada) for generously providing the cell lines used to produce conditioned media.

## Data Availability

The data generated in this study (including genomic data) are available upon request from the corresponding author.

## Ethical Approval

Collection and use of human samples and data was approved by the MUHC Research Ethics Board (protocol #2007-856). All animal studies were approved by the RI-MUHC Animal Care Committee (protocol #8081).

## Fundings

Montreal General Hospital Foundation.

Impact Grant award from the Department of Defense-Congressionally Directed Medical Research Programs, Award # CA200572.

STrOmal ReprograMing (STORMing Cancer); Cancer Research UK Grand Challenge.

## Conflict of Interest

The authors declare no potential conflicts of interest.

## Author Contributions

\# M.K, S.P, S.W contributed equally to this article as co-first author and performed most of the molecular cell biology experiments, data analysis, manuscript writing.

† V.S. and L.F. contributed equally to this article as co-last author.

L.F reviewed the data and manuscript.

L.F, V.S. and M.P. conceived the experiments, supervised the study.

N.B supervised biobank maintenance, assisted in data analysis and manuscript writing.

J.B, W.Z. and H.Z. assisted in PDOs, CAFs, and TILs generations.

R.M, H.K., B.G., K.T assisted in molecular cell biology experiments.

S.B. supervised WES analysis.

Y.Y. and N.O. assisted with WES analysis.

F.B developed PDX models and maintained corresponding biorepository.

J.T. assisted in clinical and statistical analysis.

X.S. reviewed manuscript.

G.E. and P.F. performed pathological evaluation of the samples.

## Notes

### Competing Interest Statement

The authors have declared no competing interest.

