## Supplemental Data 2 for "Optimized large-scale longitudinal biorepository of gastroesophageal adenocarcinoma patient-derived organoids: High-fidelity models for personalized treatment to overcome resistance"

**Supplementary Experimental Section**

**Initial tissue processing**

1. Tissues from consented patients are received following endoscopy or surgery.
   a. Additional biopsy samples are collected in endoscopy suite and transferred in transport media (RPMI 1640, Gentamicin (50 ug/mL), Pen/Strep (100 U/mL), Fungizone (2.5 ug/mL)).
   b. For resections, the attending pathologist identifies tumor and/or grossly normal regions, if they judge that there is tissue available for research from either or both regions, small pieces are dissected out and transferred to sterile 10 cm Petri dishes on ice.
2. Upon arrival in laboratory, wash tissue samples with sterile AD-DF+++ medium to remove excess blood and other potential contaminants.
3. Cut away and remove visibly necrotic parts of tissue (grossly white and fluffy) using a razor blade.
4. Cut the remaining tissues into the following pieces (when available):
   i. 1 small piece from tumor sample for FFPE sample
   ii. 2 fragments (1mm^3^ each) from tumor sample for implantation into mouse if PDX will be performed.
   iii. A very small piece from tumor and normal samples for TIL cultures
   iv. A moderately sized piece from tumor or normal samples for processing to a single-cell suspension and subsequent organoid and fibroblast derivation

**Tissue dissociation**

5. Mince tissues (from Step 4iv above) above using sterile razor blade and forceps in 5 mL of tissue dissociation mix (Collagen III (2 mg/mL), Hyaluronidase (100 U/mL) and Primocin (10μg/mL) in AD-DF+++ media). Collect minced tissue into a gentle MACS-C tube (Milltenyi Biotec. Cat no.130096334).
6. Process MACS-C tubes in a GentleMACS^TM^ Octo Dissociator with heaters (Miltenyi Biotec.)
   i. **Program A:** 2x30 mins for fresh surgical specimen tissue (15 min for frozen tissue). 2x15min for fresh biopsy tissue (10 min for frozen biopsy).
   ii. **Program B:** 1 min.
   iii. **Note:** At end of Program A, it is necessary to check if sample has dissociated. If not, repeat if necessary. Repeated Program A can vary depending on tissue size and state (between 15 mins and 1 hour).
7. Add 10mL of PBS + 1mM DTT to the tube and pipette up and down to mix.
8. Pass cells through 100 um cell strainers (Fisherbrand, Cat No. 22363549) into a fresh 50mL tube.
9. Transfer the sample to 15mL tube.
10. Centrifuge at 500 rcf for 5 min at 4°C.
11. Gently remove supernatant (leaving only <0.5mL behind), with pipette (not vacuum line!), resuspend pellet in 1mL of 0.25% trypsin and incubate in 37C water bath for 5 min.
12. Add 9mL PBS/10% FBS + 1mM DTT and resuspend well.
13. Divide samples into two 15mL tubes and proceed with.

**Propagation and freezing of TILs**

14. Place very small pieces from tumor regions (from Step 4iii above) into single well(s) of 24 well plate and add 500mL TILs propagating media (StemCell Cat No. 10981) with IL2 (final conc. 10^3^ CU/mL).
15. Incubate plates in Tri-gas incubator (5% CO_2_, 3% O_2_, 37°C) for TIL generation and proliferation. Do not disturb the culture and change the media only after 5-6 days.
16. If TILs leaking from tissue parts form clusters after 1-2 weeks, collect the cells in 15mL Falcon Tubes and centrifuge at 500 rcf for 5 min.
17. Discard supernatant and break the pellets by adding fresh TILs media. Passage as above for 3 generations before freezing as viable cells (Step 18).
18. Freeze cells slowly (1C/min) in 10% DMSO/FBS mixture. Store in liquid nitrogen tank.

**Propagation of organoids and fibroblasts**

19. If proceeding with scRNAseq/Protocol 5), take 1mL of solution from 15mL tube from step 12 above (10mL cells + PBS + 10% FBS) and transfer to a new 15mL conical tube.
    a. If not proceeding with Protocol 5, use the original 15mL tube from Step 12 above and proceed to Step 20 below.
20. Centrifuge 15mL tube at 500 rcf for 5 min at 4 degrees.
21. Remove supernatant and resuspend in 4mL AD-DF+++. Let stand for 10min, then transfer 2mL of the upper part of the supernatant to a new 15mL conical tube (this portion is used for fibroblast culture). Centrifuge both fractions at 500 rcf for 5 min at 4 degrees.
22. **For fibroblast propagation**: Remove supernatant, add 1mL fibroblast medium, resuspend the cell pellet and transfer into Type I collagen precoated 24 well plate.
23. **For organoid propagation:** Remove as much supernatant as possible with p1000 pipette tip. Spin down quickly and remove as much supernatant as possible using a p200 pipette.
    a. Place the tube on ice to cool down for a few minutes. Add 80uL Matrigel to cell pellets and mix well.
    b. Add cell + Matrigel mix into a new pre-warmed 24-well plate.
    c. Allow the plate to incubate for 1-2 min at RT, before transferring it to the 37°C/5% CO_2_/3% O_2_ Tri-gas incubator for 5-10 mins to allow the Matrigel to become a solid dome.
    d. Add 600uL of GEA PDO propagation media (1:1 mix of IntestiCult and conditioned medium) to each well of 24 well plate.
    e. Return plate to Tri-gas incubator

**Sample preparation for single-cell RNA sequencing (scRNA seq)**

24. Centrifuge 15mL tube containing 9mL solution (from step 13 above) at 500 rcf for 5min at 4°C.
25. Remove supernatant with 5mL pipette, resuspend pellet in 0.5mL Dispase and 10uL DNase I (1 mg/mL), and incubate in 37°C water bath for 5 min.
26. Add 9mL PBS.
27. Centrifuge at 500 rcf for 5 min.
28. Aspirate supernatant with pipette (leave some PBS behind), not vacuum, resuspend pellet in 1mL ACK lysis buffer (Invitrogen Cat No. A1049201) for 5 mins at RT.
29. Add 9mL 1X PBS.
30. Pass cells through a 40 um cell strainer (Falcon, Cat No. 352340).
31. Centrifuge at 500 rcf for 5 mins.
32. Aspirate supernatant with a pipette.
33. Resuspend in 1mL 2% FBS-PBS and transfer in a 1.5mL microtube.
34. Centrifuge at 500 rcf for 5 min.
35. Wash twice following steps (10-11).
36. Resuspend in 100uL 2% FBS-PBS for further processing.
37. Sample is now ready for testing of viability, RBC content and debris at Genome Center for decision-making on whether or not to move ahead with library construction)
    a. **Note:** Normal tissue yields fewer cells than tumor tissue for an equivalently sized input sample.

**PDXO generation**

38. Previous steps – perform PDX generation and tissue retrieval as per established procedures.
39. Bring excised PDX in a dish with PBS to BSC in the lab.
40. Wash PDX with fresh sterile 1XPBS.
41. Using forceps and a razor, mince the tumor until it is almost devoid of visible tumor mass.
42. Transfer the tumor to a Miltenyi C-tubes containing 5mL Tissue dissociation mix
43. Incubate C-tubes on the gentle-MACS Octo Dissociation counter. Dissociate tissues at 37°C for 30min (as per Step 6 above).
44. Bring dissociated sample back to BSC and add 10mL of PBS.
45. Filter tissue suspension through a 100μm strainer into a 50mL tube.
46. Transfer contents to a 15mL tube and centrifuge at 500 rcf for 4 min.
47. Remove supernatant without disturbing the pellet.
    a. **Note:** Use a pipette if small pellets are left.
48. If there are visible red blood cells, add 1-2mL of ACK Lysis Buffer to the pellet, pipette up and down and incubate for 3 mins at RT.
49. Add 10mL of PBS and centrifuge at 500 rcf for 4 mins.
50. Remove supernatant and add 1mL 1X PBS.
51. Perform a cell count (before proceeding with mouse cell depletion). Measure single cell viability using Trypan Blue assay.
    a. Mix 10uL of Trypan Blue solution with 10uL of cell suspension. Count cell numbers with Countess II FL (Invitrogen).
52. Centrifuge cell suspension at 300×g for 10 mins. Aspirate supernatant completely.
53. Resuspend cell pellets in AD-DF+++ medium containing mouse depletion antibody cocktail as per manufacturer instructions (Miltenyi Biotech. cat no.130-104-694). Briefly, 80uL of cell suspension (AD-DF+++ medium) with 20uL of mouse depletion antibody cocktail per 2×106 tumor cells or 107 total cells (including red blood cells). Pipette up and down using a P1000 pipette and incubate in refrigerator (2-8°C) for 15 minutes.
54. Organize MACS: Multistrand (MACS Separators) apparatus and MACS LS Columns.
55. In the hood, set up the stand and attach the filter onto the magnetic portion.
56. Add 3mL of AD-DF+++ medium to the magnetic column to wash and calibrate before use.
57. Add 500uL of AD-DF+++ medium to the mixture and mix well.
58. Pass the cell suspension through the column and collect the flowthrough in a new sterile 15mL tube.
59. Pass another fresh 1mL of AD-DF+++ medium through the column.
60. If doing another sample, replace the filter, and repeat the steps above for each sample.
61. Determine the cell count and cell viability of both cell fractions (pre- and post-mouse cell depletion). Follow the protocol from Step 51.
62. Centrifuge collected (mouse cell-depleted fraction) cells at 500 rcf for 4 min.
63. Prepare 24 well-plates with complete organoid propagation media (supplementary) + 5% Matrigel.
64. Discard media and resuspend the cell pellet with complete organoid propagation media. Add this to the well and incubate in 37°C Tri-gas incubator.
    a. **Note:** PDXO propagation requires complete organoid propagation media with 5% Matrigel.

**Passaging and freezing organoids**

65. Prepare Matrigel digestion buffer: 1:100 dilution of 100 mg/mL stock Collagenase/Dispase (Collagenase/Dispase 100 mg/mL (Sigma 10269638001) with AD-DF+++ medium.
66. Remove existing medium in organoid wells and add 500uL digestion buffer in each well
67. Pipet up and down to break the Matrigel domes.
68. Digest for about 1h at 37°C.
69. Collect well contents in a 15mL tube.
70. Spin at 500 rcf, 5 min.
71. Carefully remove supernatant, add 1mL of 0.25% Trypsin.
72. Incubate at 37°C for 3 x 5 min. After each incubation, vortex vigorously to break up the organoids.
73. Add 4mL of 5% FBS-PBS to inactivate the trypsin. Vortex to generate a smooth suspension of cells.
74. Spin at 500 rcf, 5 min.
75. Remove supernatant, add 1mL AD-DF+++ medium to wash.
76. Count cells (see Step 51). Plate 30K per new well for ongoing propagation and slow-freeze (1C/min) the rest of the cells in CryoStor CS10 (StemCells, Cat No. 07930)
77. Plating in new wells:
    a. Spin down at 500 rcf for 5 min at 4°C. Carefully pipet off and discard the supernatant.
    b. Do an extra quick spin and remove all the media with a smaller pipette tip.
    c. Place the tube on ice for 3 minutes.
    d. For one well, resuspend 35,000 cells in 80uL Matrigel, on ice. In a 24w plate, plate the Matrigel + cells delicately so it forms a dome.
    e. Allow the plate to incubate for 1 min at RT, before transferring it to the 37°C in 3% O2 incubator for 10-20 mins to allow the Matrigel to become a solid dome.
    f. Add 600uL of complete organoid growth medium to each well of 24 well plate.
    g. Refresh media 3-4 days. Passage every 10-14 days depending on the growth of organoids.
    h. **Notes:**
       i. Always keep Matrigel on ice - the evening before procedure, remove Matrigel from -80°C freezer and leave to thaw on ice in 4C fridge overnight.
       ii. Organoids (PDOs) must be cultured in 100% Matrigel.

**Passaging fibroblasts**

78. Fibroblasts adhere at the bottom of collagen-coated plate (see Step 22)
    a. Coating protocol: Dilute Purecol (collagen I) solution (1:30) in DPBS. Use 400uL per well (24-well plate), 1mL per well (6w plate) or 5mL per plate (10 cm petri dish). Incubate at least 30 min, ideally 2h, at 37°C. Remove Purecol, wash twice with 5mL DPBS, aspirate all the liquid and store plates at 4C for up to 2 weeks.
79. When cell confluency reaches 80-90% (or if fibroblasts are growing in patches) discard the media.
80. Wash cells once with 1mL 1X DBPS. Aspirate DPBS and add 500uL 0.25% trypsin. Incubate 3-5 minutes at 37°C.
81. Add 1mL trypsin inactivating solution (5% FBS in 1X DPBS) into the wells. Pipet the cells up and down to detach the rest of adhered cells and transfer them in a 15mL tube.
82. Centrifuge 5 min, 500 rcf, then carefully remove supernatant.
83. Resuspend the pellet in 1mL Fibroblast media.
84. If the 24w plate was confluent, transfer all the cells to a pre-coated 10 cm petri dish (cells are now at passage 2 (or P2)); if you had patches of cells, it is better to transfer to a 6-well pre-coated plate. When the 6-well plate is confluent, detach the cells and transfer them to a 10 cm petri dish (cells are then considered to be at passage 3/P3).
85. Once 80-90% confluency is achieved, freeze fibroblasts following the protocol mentioned below.
    a. Remove media, wash with 5mL DPBS.
    b. Add 5 mL 0.25% trypsin, incubate 3-5 min at 37°C.
    c. Add the 5mL 5% FBS-DPBS to inactivated trypsin.
    d. Pipet out the cells-mix in a 15mL tube, centrifuge 5 min at 500 rcf.
    e. Remove supernatant.
    f. Label 3 cryovials (biobank code, cell type, passage, date).
    g. Resuspend the pellet in 1.5mL **Freezing solution (**90% FBS-10%DMSO).
    h. Distribute 0.5mL in each cryovial. Place the vials at Mr-Frosty freezing containers (Thermofisher, cat no. 5100-0036) and place them at -80°C for at least 4 h before transferring them to Liquid N_2_.
    i. **Note:** Fibroblasts stop growing if they get overcrowded.

**PDXO passaging and freezing**

86. Take PDXO plates from the 37°C Tri-gas incubator to BSC.
87. Discard media from plates.
88. Add 500uL Matrigel digestion buffer (step 65) at each well.
    a. Break the Matrigel layer containing matured organoids using pipette. Let plates sit in 37°C incubator for at least 45 minutes.
89. Collect cells in labelled 15mL tubes.
90. Centrifuge tube at 500 rcf for 5 min in the culture room centrifuge.
91. Discard the supernatant as much as possible without disrupting the pellet.
    a. **Note:** If there is no pellet, discard the supernatant first, then use pipette to take away the rest, leaving the cloudy part (as this may still have organoids)
92. Add 1mL of trypsin to the pellet, pipette up and down and incubate in the 37°C water bath for 10-15 min.
93. Every 5 min check on cells. Flick the tube, pipette up and down, and check under microscope to see if they are dissociated into single cells. If not, incubate for longer.
94. Once cells appear to be single, add DMEM+10%FBS to trypsin tube to complete to 14mL.
95. Centrifuge tube at 500 rcf for 5 min.
96. Remove the supernatant and then pipet out all liquid possible without disrupting the pellet.
97. Add 2mL of organoid medium and mix.
98. Determine cell count and cell viability (using Trypan Blue).
99. In a 15mL tube add complete organoid media with 5% Matrigel.

100. Add 1mL of mixture to each coated plate and incubate in 37°C tri-gas incubator.

**Flow cytometry for characterization of patient-derived cells**

PDO, TILs or fibroblasts were resuspended in FACS buffer (1x PBS, 2-5 % (v/v) FBS (or BSA, 0.5 mM EDTA, 2 mM NaN3). After centrifugation of samples at 300g for 5 min at 4°C, 100uL of Aqua Blue 405nm (viability control, 1:1000 dilution in 1X PBS) was added to the cell pellet and samples were incubated for 30 min on ice in the dark. Afterwards, 500uL PBS was added to each tube to wash the dye, and then samples were spun down at 300g for 5 min at 4°C. After removal of supernatant, 100uL of appropriate antibody was added to each sample and samples were incubated for 30 min on ice in the dark, followed by PBS washing and centrifugation at 300g for 5 min at 4°C. Lastly, the stained cell pellets were resuspended in 300uL FACS buffer per tube and kept on ice until ready for FACS assay.

**Immunofluorescence staining of fibroblasts**

PureCol collagen was used to coat coverslips, followed by plating of fibroblasts. When cells reached 70-80% confluent, media was removed and cells were washed with PBS, followed by fixation in 4% paraformaldehyde (PFA) for 10 min at room temperature and permeabilization in 0.1% Triton X in PBS for 10 minutes at room temperature. Immunofluorescence staining was carried out using manufacturer’s suggested dilutions of the primary, secondary antibodies and DAPI nuclear stain. Stained fibroblasts were imaged using a Zeiss LSM 780 confocal microscope. Primary antibodies used are Phalloidin (ThermoFisher, A22287), 𝒶SMA(ThermoFisher, MA5-11547), PDGFRβ (R&D system, AF385-SP). Secondary antibodies used were donkey anti-goat Alexa Fluor 488 (Invitrogen, A11055) and goat anti-rabbit Alexa Fluor 555 (Invitrogen, A31572).

**Supplementary Tables**

**Supplementary Table 1:** **Tab 1**, “Patient Cohort”, lists of epidemiological, clinical and pathological variables and PDMs generated for each of the 382 patients for whom this was attempted. NS, normal stomach; T, tumor; Tsur, tumor from surgery; NE, normal esophagus; TNR, treatment-naïve resection. **Tab 2**, “by stage”, PDM success rate by tumor stage. Tbio, tumor from biopsy; Tsur, tumor from surgery. **Tab 3**, “by grade”, PDM success rate by histological grade. **Tab 4**, “by sex”, PDM success rate by patient sex. **Tab 5**, “by MMR status”, PDM success by MMR status. MSI-H, microsatellite instability-high; MSS, Microsatellite-stable. **Tab 6**, “by treatment response”, PDM success rate by pathological response to therapy (Tumor Regression Grade, TRG; TRG 0, complete response; TRG 1, near-complete response; TRG 2, partial response; TRG 3, minimal or no response).

**Supplementary Table 2:** Expansion media for GEA PDO propagation (EM)

| **Component** | **Final concentrations** |
| --- | --- |
| Conditioned medium (Cell line: HEK293 (Wnt3a, R-Spondin, Noggin Recombinant)) | 50%vol/vol |
| IntestiCult Medium (StemCell Technologies, 06010) | 50% vol/vol |
| GlutaMAX (Thermo Fisher Scientific, 35050-061) | 1x |
| HEPES (Thermo Fisher Scientific, 15630-106) | 10mM |
| B27 (Thermo Fisher Scientific, 17504-044) | 1x |
| N2 (Thermo Fisher Scientific, 17502-508) | 1x |
| Nicotinamide (Sigma-Aldrich, N0636) | 10mM |
| NAC (N-acetyl-l-cystein) (Sigma-Aldrich, A5099) | 1.25mM |
| mEGF (Peprotech, 315-09) | 50ng/mL |
| hFGF10 (Peprotech, 100-26) | 200ng/mL |
| LeuGastrin (Sigma-Aldrich, G9145) | 10nM |
| A8301 (Tocris, 2939) | 500nM |
| SB202190 (Sigma-Aldrich, S7067) | 10uM |

**Supplementary Table 3:** Other PDM propagation media

| **Fibroblast Medium** |  |
| --- | --- |
| **Component** | **Final Concentration** |
| DMEM+Glutamax-1 (Invitrogen 10569-010) |  |
| Gentamicin (Invitrogen 15710-064) (10mg/mL) | 50ug/mL (1:200) |
| EGF (100ug/mL) | 5ng/mL (1:20,000) |
| Insulin (Invitrogen 12585-014) (4mg/mL) | 5ug/mL (1:800) |
| BPE (Wisent 002-011-IL) (5mL) | (1:400) |
| Normocin (Invivogen ANT-NR2) (50mg/mL) | 100ug/mL (1:500) |
| FBS | 10% |
| **Transport medium** |  |
| **Component** | **Final Concentration** |
| RPMI 1640 (Invitrogen 11875-093) |  |
| Gentamicin (Invitrogen 15710-064) (10mg/mL) | 50ug/mL (1:200) |
| Pen/Strep (Sigma P4333) (100x) | 100U/mL |
| Fungizone (Invitrogen 15290-026) (250ug/mL) | 2.5ug/mL (1:100) |
| **AD-DF+++ medium** |  |
| **Component** | **Final Concentration** |
| Advanced DMEM/F12 (Invitrogen 12634-010) |  |
| Glutamax-1 (Invitrogen 35050-061) (200mM) | 2mM (1:100) |
| HEPES (Invitrogen 15630-080) (1M) | 10mM (1:100) |
| Pen/Strep (Sigma P4333) (100x) | 100U/mL |
| **IntestiCult medium** |  |
| **Component** | **Final Concentration** |
| Component A (Stemcell #100-0191) |  |
| Component B (Stemcell #100-0190) |  |
| Pen/Strep (Sigma P4333) (100x) | 100U/mL |
| ROCK inhibitor (50mM) | 10uM (1:5000) |
| **Immunocult medium** |  |
| **Component** | **Final Concentration** |
| Immunocult (StemCell XF #10981) |  |
| IL-2 (StemCell #78036) | 10^3^CU/mL |
| Pen/Strep (Sigma P4333) (100x) | 100U/mL |
| Normocin (Invivogen ANT-NR2) (50mg/mL) | 100ug/mL (1:500) |
| **Tissue dissociation mix** | **For 5mL** |
| Collagen III (Cedarlane Labs LS004182) | 500uL |
| Hyaluronidase | 50uL |
| Primocin | 10uL |
| AD-DF+++ media | 4.5mL |

**Supplementary Table 4:** Clinical information on 22 drugs shown in the chemogram.

| **Drug name** | **Route of administration** | **Target** | **FDA approval** | **Clinical trial for**  **ESO/GEJ/GAS cancer** | **Clinical trial reference** |
| --- | --- | --- | --- | --- | --- |
| Idarubicin | IV | Topoisomerase | YES | NO | – |
| Vincristine | IV | Microtubule | YES | YES | Rake et al., 1979 (GAS) |
| Vinblastine | IV | Microtubule | YES | NO | – |
| Gemcitabine | IV | Nucleoside Analog | YES | YES | NCT04336098 (GEJ) |
| Daunorubicin | IV | Topoisomerase | YES | NO | – |
| Topotecan | IV and PO | Topoisomerase | YES | YES | NCT00252889 (ESO/GAS) |
| Paclitaxel | IV | Microtubule | YES | YES | NCT03550482 (GAS) |
| Mitoxantrone | IV | Topoisomerase | YES | YES | NCT04718402 (GAS) |
| Docetaxel | IV | Microtubule | YES | YES | NCT00849615 (GEJ) |
| Ixabepilone | IV | Microtubule | YES | YES | NCT00017043 (GEJ) |
| Epirubicin | IV | Topoisomerase | YES | YES | NCT01726452 (GEJ) |
| Vinorelbine | IV and PO | Microtubule | YES | YES | NCT00215462 (ESO/GAS) |
| Pirarubicin | IV | Topoisomerase | NO | YES | Kikuyama et al., 1998 (GAS) |
| Ixazomib | PO | Proteasome | YES | NO | – |
| Doxorubicin | IV | Topoisomerase | YES | YES | NCT00165191 (ESO/GAS) |
| Cabazitaxel | IV | Microtubule | YES | YES | NCT01956149 (GEJ) |
| Teniposide | IV | Topoisomerase | YES | YES | Berenberg et al., 1993 (GAS) |
| Cytarabine | IV and SC | Nucleoside Analog | YES | NO | – |
| Pralatrexate | IV | Antifolate | YES | YES | NCT01178944 (GEJ) |
| Raltitrexed | IV | Nucleoside Analog | NO | YES | NCT03083613 (GEJ) |
| 5-Fluorouracil | IV | Nucleoside Analog | YES | YES | NCT00849615 (GEJ) |
| Oxaliplatin | IV | DNA base | YES | YES | NCT00849615 (GEJ) |
