## Supplemental Data 1 for "Optimized large-scale longitudinal biorepository of gastroesophageal adenocarcinoma patient-derived organoids: High-fidelity models for personalized treatment to overcome resistance"

### Supplementary Figure 1

A.

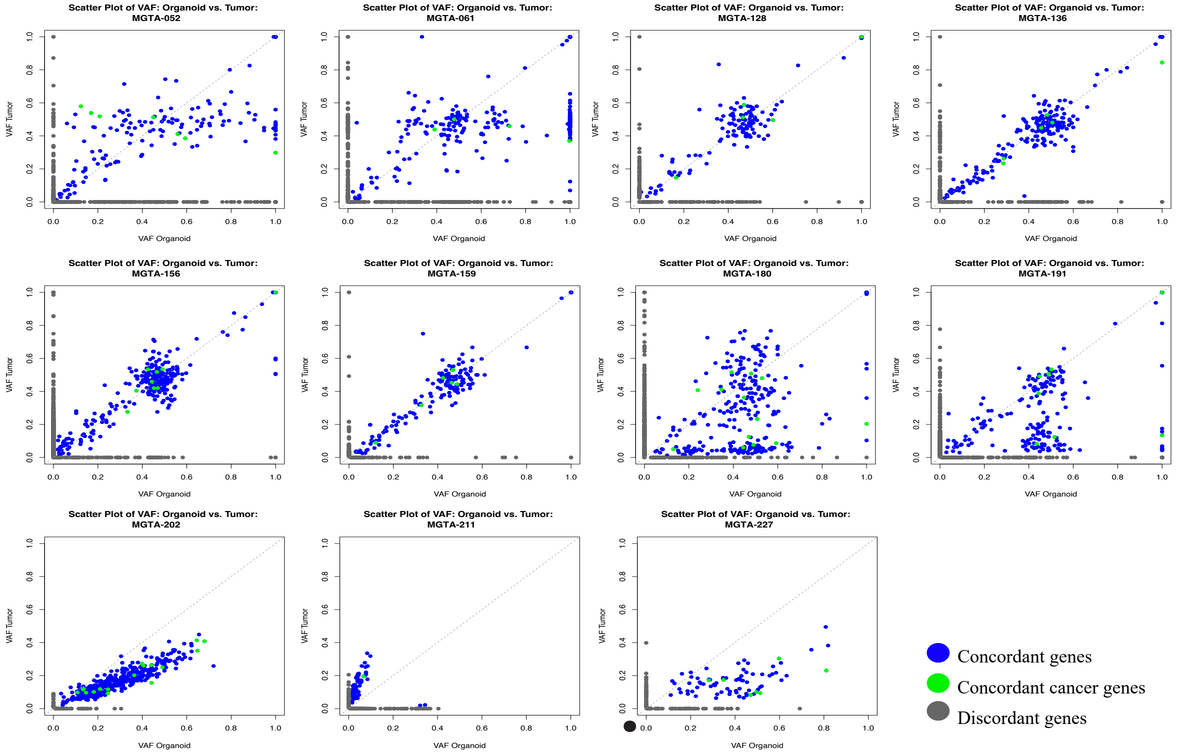


B.

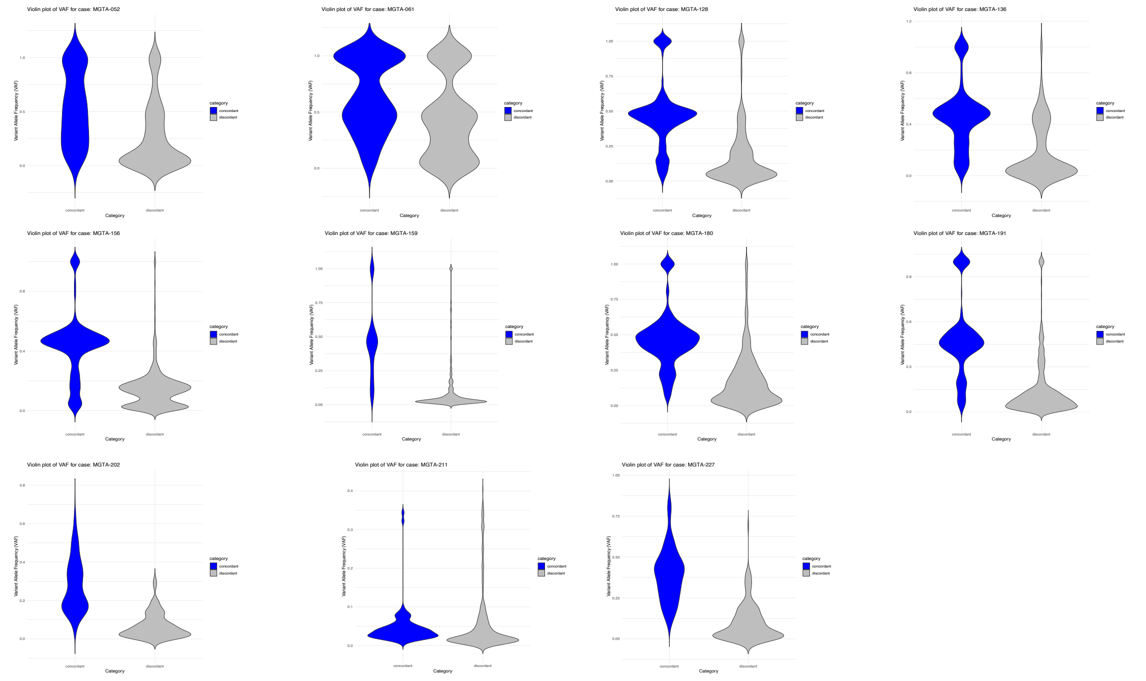


**Supplementary Figure 1. Genomic profile comparison between PDO and matched primary tumors across 11 pairs.**

**A.** Scatter plots showing the correlation of VAFs between PDOs and their matched tumors. Each dot represents a unique point mutation. Blue dots are concordant mutations between tumor and organoid; green dots are concordant cancer gene mutations; grey dots are discordant mutations. The dashed line represents the identity line (x = y). **B.**Violin plots showing the distribution of VAFs for concordant (blue) and discordant (grey) mutations across the same 11 PDO-tumor pairs, shown in the same order.

### Supplementary Figure 2

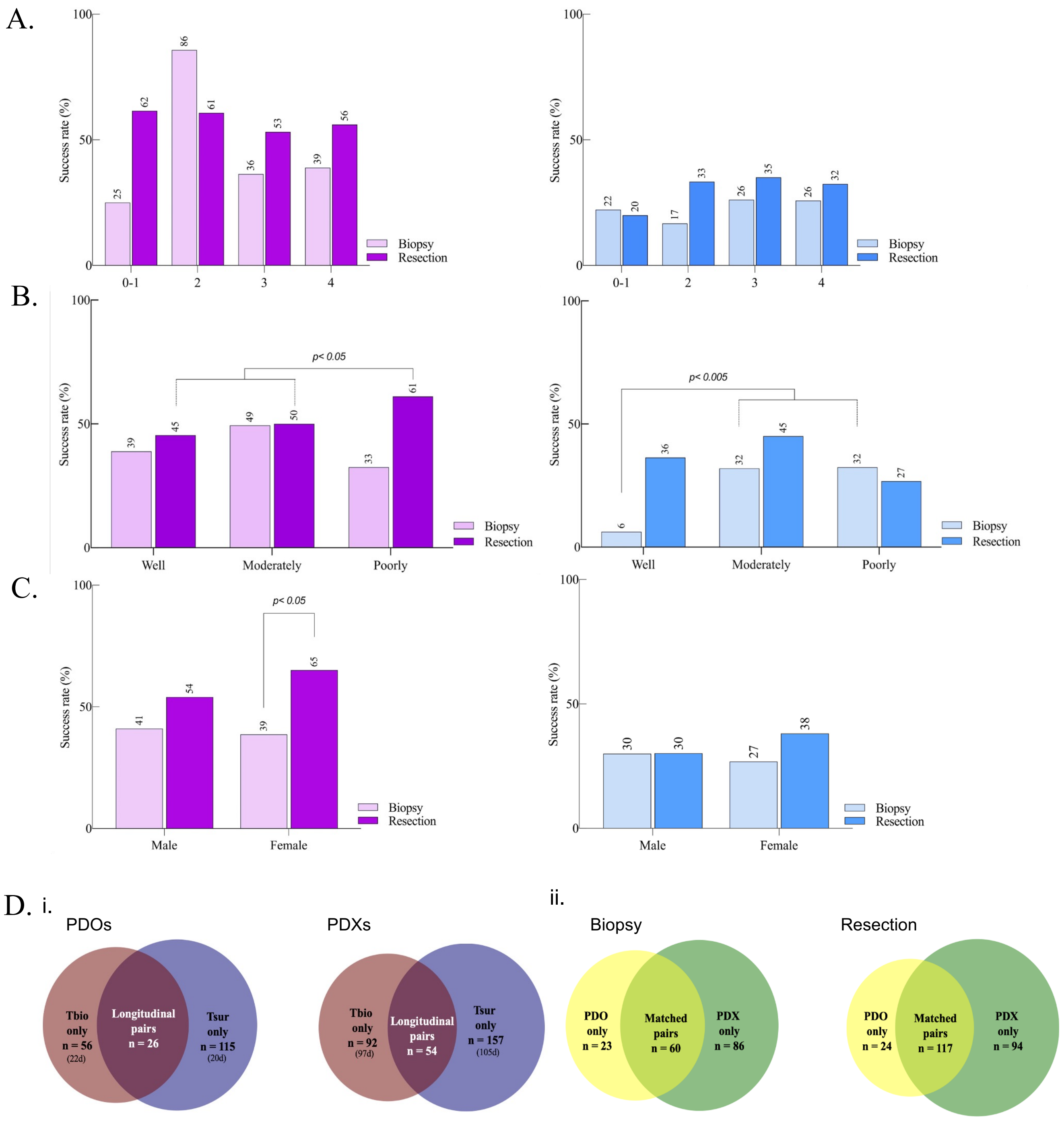


**Supplementary Figure 2. Determinants of successful PDM generation. A.** Success rate for samples separated by clinical stage for PDO (left) and PDX (right) generation. **B.** Success rate for samples separated by histopathological grade for PDO (left) and PDX (right) generation. **C.** Success rate for samples separated by sex for PDO (left) and PDX (right) generation. Only p values with significance are shown in the figure. **D.** i. The Venn diagrams display the number of patients for whom PDOs (left) and PDXs (right) were successfully generated from samples taken during biopsies and resections (either treatment-naïve resection group or received NACT). The overlapping section indicates the number of patients for whom longitudinal PDOs or PDXs were created, encompassing both pre- and post-treatment stages. ii. Additional Venn diagrams depict the exclusivity as well as overlaps in the generation of PDOs versus PDXs for samples derived from biopsies (left) and resections (right) from the same patients.

### Supplementary Figure 3

A. Epithelial lineage of PDOs

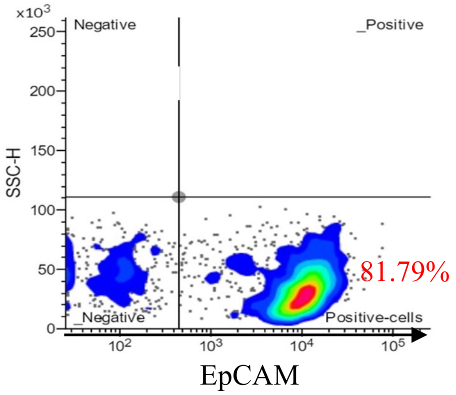


B. Human epithelial lineage of PDXOs

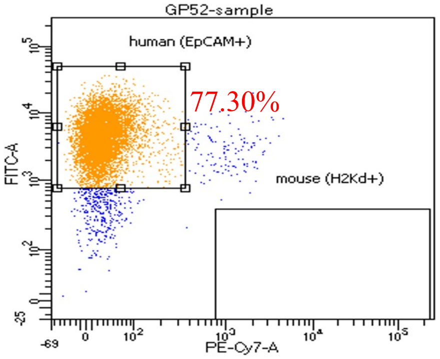


C. Stromal lineage of CAFs

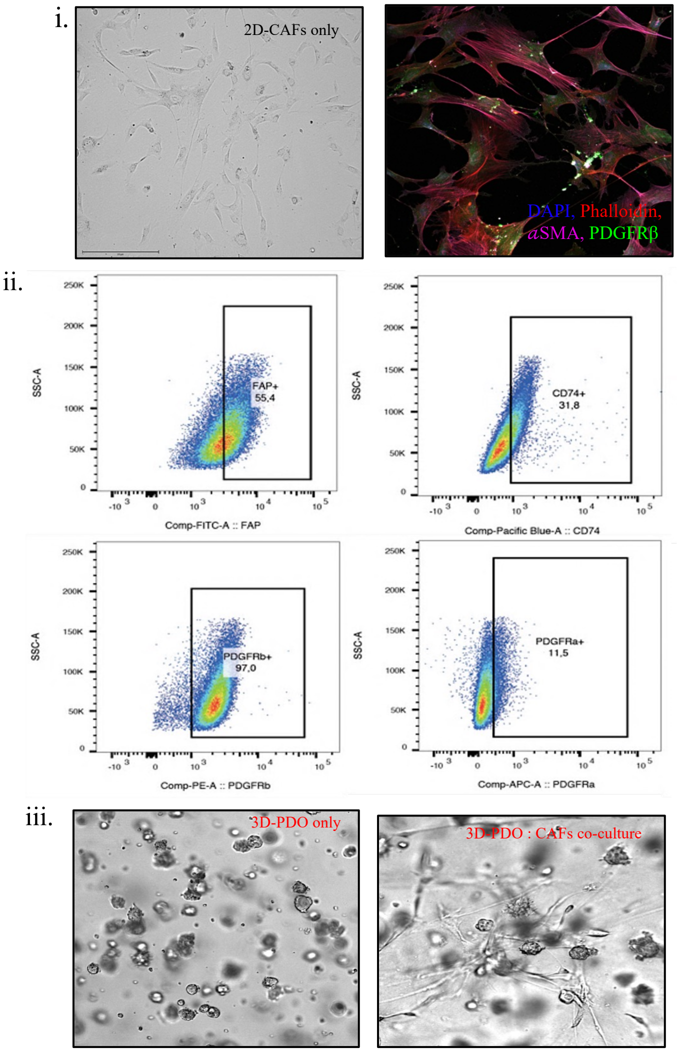


D. Immune lineage of TILs

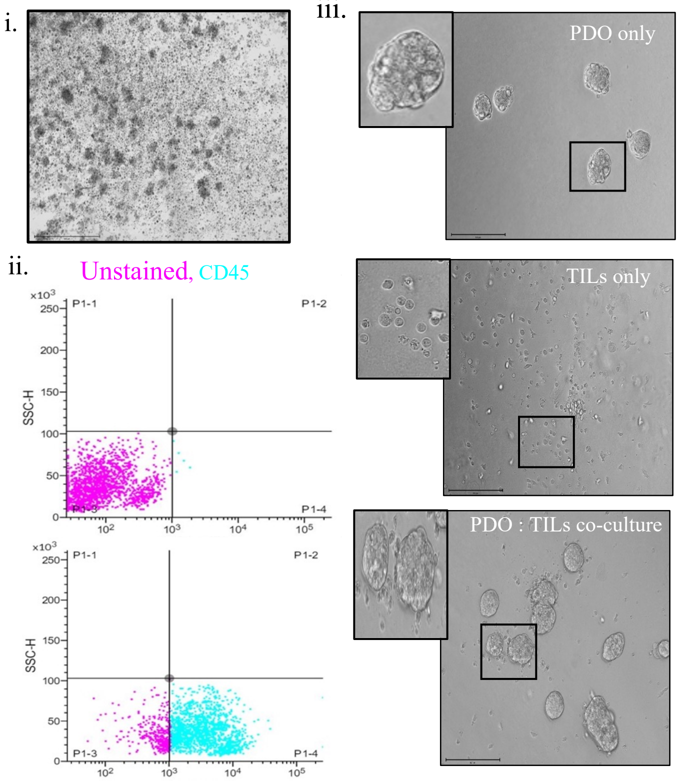


**Supplementary Figure 3. Purity and quality of cells derived from PDOs and stromal fractions (CAFs and TILs) was confirmed through different methods A.** FACS analysis of a representative PDO sample demonstrates that most cells contained therein are of epithelial origin. **B.** FACS analysis of a representative PDX sample demonstrates that most cells contained therein are of human origin. **C.i.** Isolated CAFs display expression of CAF-specific markers as visualized by IF (phalloidin, *α*SMA, PDGFRβ) and **ii.** FACS (FAP, CD74, PDGFRβ, PDGFR*α*). **iii.** CAF-PDO co-culture model for evaluating CAF-PDO interplay for chemoresistance. **D i.** Bright-field image of a representative patient-derived TIL 2D culture. **ii**. Isolated TILs express the hematopoietic lineage-specific maker CD45 as assessed by FACS. **iii.** A High-throughput PDOs and TILs co-culture model for testing the efficacy of drugs precisely immunotherapy.

### Supplementary Figure 4

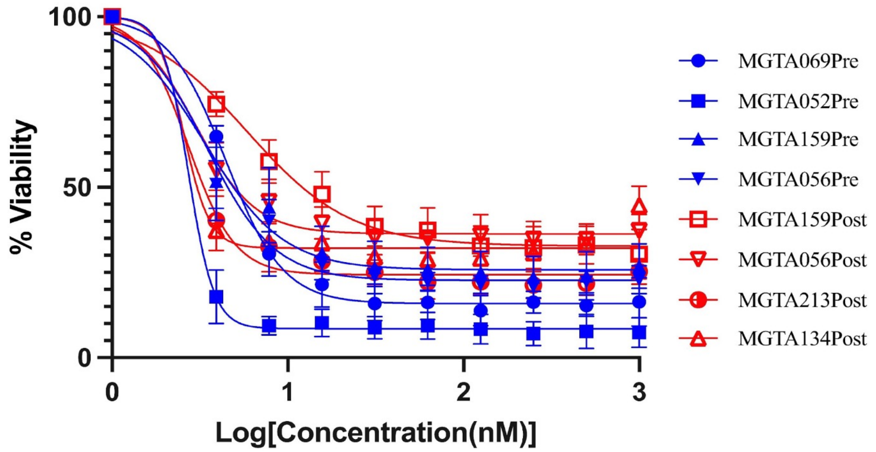


| | Pre | Post |
|---|---|---|
| Normalized AUC | 0.41 | 0.55 |

**Supplementary Figure 4. PDOs recapitulate longitudinal chemotherapy response.** Survival curves showing the response of four biopsy-derived (blue, Pre) and four resection-derived (red, Post) PDOs to DCF treatment. PDOs from resected tissues exhibit higher resistance, as reflected by increased Normalized AUC value, indicating enhanced chemo-resistance following treatment.
